# Single-cell in situ transcriptomic map of astrocyte cortical layer diversity

**DOI:** 10.1101/432104

**Authors:** Omer Ali Bayraktar, Theresa Bartels, Damon Polioudakis, Staffan Holmqvist, Lucile Ben Haim, Adam M.H. Young, Kirti Prakash, Alexander Brown, Mercedes F. Paredes, Riki Kawaguchi, John Stockley, Khalida Sabeur, Sandra M. Chang, Eric Huang, Peter Hutchinson, Erik M. Ullian, Daniel H. Geschwind, Giovanni Coppola, David H. Rowitch

## Abstract

During organogenesis, patterns and gradients of gene expression underlie organization and diversified cell specification to generate complex tissue architecture. While the cerebral cortex is organized into six excitatory neuronal layers, it is unclear whether glial cells are diversified to mimic neuronal laminae or show distinct layering. To determine the molecular architecture of the mammalian cortex, we developed a high content pipeline that can quantify single-cell gene expression *in situ*. The Large-area Spatial Transcriptomic (LaST) map confirmed expected cortical neuron layer organization and also revealed a novel neuronal identity signature. Screening 46 candidate genes for astrocyte diversity across the cortex, we identified grey matter superficial, mid and deep astrocyte identities in gradient layer patterns that were distinct from neurons. Astrocyte layers formed in early postnatal cortex and mostly persisted in adult mouse and human cortex. Mutations that shifted neuronal post-mitotic identity or organization were sufficient to alter glial layering, indicating an instructive role for neuronal cues. In normal mouse cortex, astrocyte layer patterns showed area diversity between functionally distinct cortical regions. These findings indicate that excitatory neurons and astrocytes cells are organized into distinct lineage-associated laminae, which give rise to higher order neuroglial complexity of cortical architecture.

## Introduction

Cortical structure has classically been described in terms of six excitatory neuronal layers, generated sequentially during early development (*1*). In addition, the cortex is regionally specialized into areas responsible for motor, sensory and cognitive functions (*1*). Although neurons have are known to be a diverse population, astrocytes, which comprise about 50% of brain cells, are generally regarded as relatively functionally homogeneous and interchangeable. Might glia also be regionally specialized to confer added cortical organizational complexity? Gray matter astrocytes generally support central nervous system (CNS) structure, nutrition, regulation of blood flow, the blood brain barrier, synapse formation and activity (*2*). While several studies show functional diversity of astrocytes (*3–7*) with implications for regional neural circuit activity or survival (*8*), precise molecular features of glial spatial organization underlying cortical architecture remain unclear.

Single-cell spatial transcriptomic approaches provide unprecedented detail on cellular diversity (*9*), and have been used to investigate the spatial organization of the mammalian brain (*10–12*). While some approaches produce highly multiplexed gene expression information, imaging and high-content data analysis restrict their application to relatively small tissue areas. Thus, they leave open the important question of how spatial transcriptomics can be applied to screen large tissue areas, including archival tissues and human organs, in a single-cell and quantitative manner to demonstrate gene expression gradients across regions.

Here we developed a method for automated quantitative high-content screening of *in situ* single-cell gene expression in mammalian brain. We screened 50 candidate genes for astrocyte spatial organization against known patterns for neurons comprehensively in the cerebral cortex. Surprisingly, while astrocytes expression patterns showed laminar and area organization such glial layers did not correspond to the six excitatory neuronal layers, indicating higher order complexity of cortical architecture.

## Results

### Automated pipeline to map single cell cortical gene expression *in situ*

To map single cell gene expression across large tissue areas at scale, we automated multiplexed branched-DNA single-molecule fluorescent in situ hybridization (RNAScope smFISH, Advanced Cell Diagnostics) with favorable signal-to-noise ratio (*13*), immunohistochemistry (IHC), fast confocal imaging, cellular segmentation and quantification on standard whole mouse brain cryo-sections (Fig 1A). We used spinning disk confocal microscopy, to image whole tissue sections at high XY and Z resolution, then modified a high-content screening workflow software (Harmony, Perkin Elmer) to quantify RNA transcripts per cell. We manually segmented cells into brain regions based on reference atlases for gene expression analysis (see Extended Methods) to generate quantitative 2D tissue maps showing spatial single cell regulation of gene expression.

**Figure 1:**
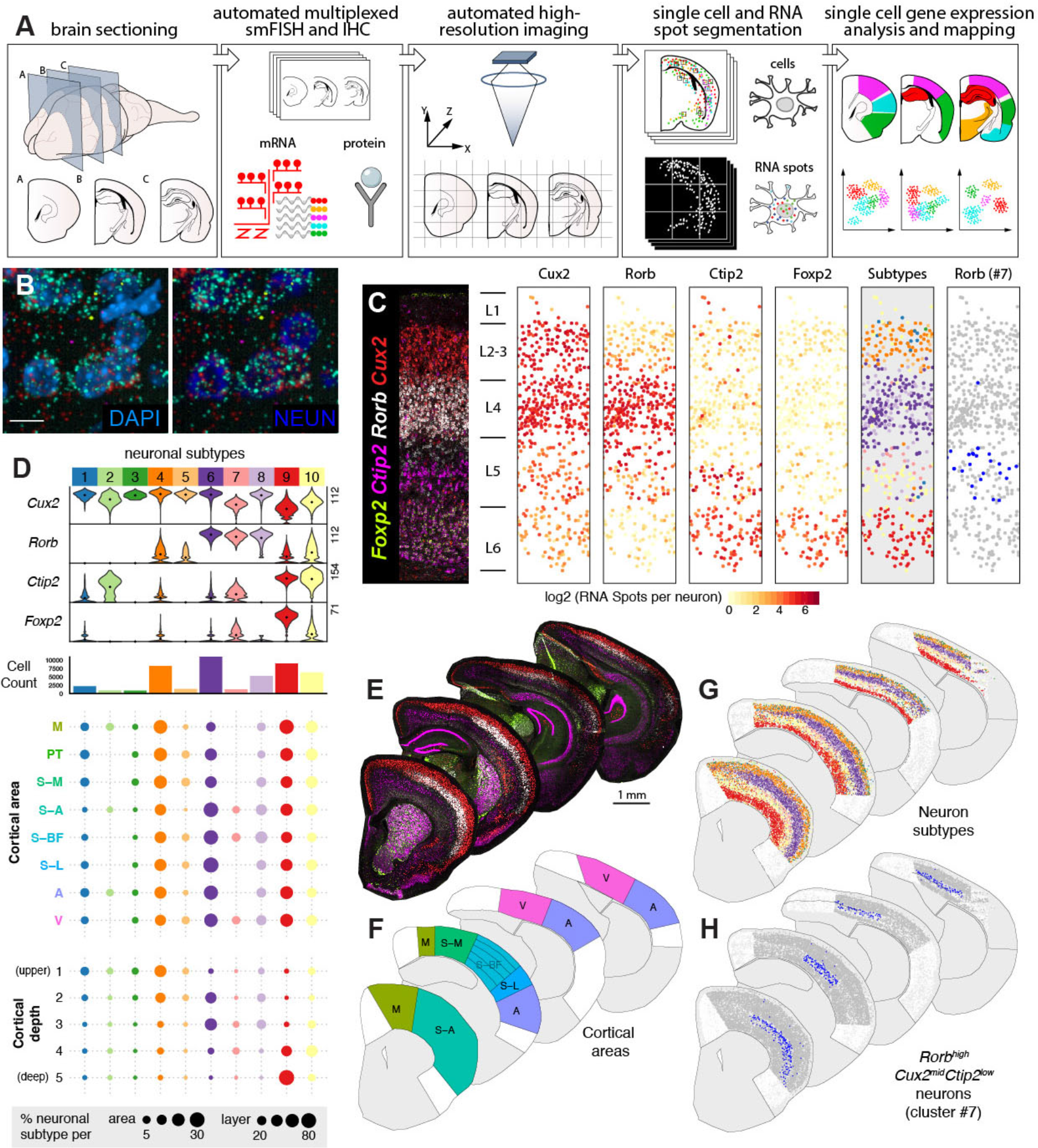
LaST map pipeline for mapping cortical neuronal subtypes *in situ*. **A)** Design of automated spatial transcriptomic pipeline. **B)** Large tissue areas are imaged entirely at high resolution to map single cell gene expression. Shown are 40X z-projection images of *Rorb*^+^ L4 neurons in the P14 mouse barrel cortex. Neuronal soma are marked by NEUN IHC and nuclei are marked by DAPI. Single RNA molecules for 4 layer neuronal markers, indicated in (E), are visualized as submicron sized puncta and quantified per cell. **C)** Automated mapping of layer neuron marker expression and layer neuron subtypes. Left panel shows a stitched image across the depth of the mouse barrel cortex. Middle panels show maps of automatically identified single neurons, plotted as solid circles and colored quantitively for RNA spot counts per gene per cell. Right panels show the spatial distribution of neuronal subtypes identified in (D). **D)** Identification of neuronal subtypes based on unbiased classification of single cell level smFISH data. 10 major neuronal populations were identified based on tSNE and hierarchical clustering of 46,888 neurons across 8 broad cortical areas for the expression of 4 layer neuron markers. (Top) Violin plots show single cell expression profiles of clusters, highest RNA spot count per cell are shown on the left. (Middle) Histograms showing total number of cells per cluster. (Bottom) Spatial distribution of clusters across cortical areas and five normalized cortical depth bins, shown as percentage of total neurons in given area/depth bin (bottom). **E-H)** Single cell mapping of cortical neuron subtypes: (E) Low magnification images of P14 hemisections from four different anatomical levels, (F) broad cortical areas included in the analysis, (G) maps of 10 major neuronal populations, and (H) spatial distribution of area-restricted L5 *Rorb*^*high*^*Cux*^*mid*^*Ctip2*^*low*^ neurons. Scalebars: (B) 10 μm, (E) 1 mm. Abbreviations: M, motor, S-A, anterior somatosensory, S-M, medial-somatosensory, S BF, somatosensory barrel, S-L, somatosensory-lateral, PT, parietal, A, auditory, V, visual.

### LaST map confirms neuron excitatory layering and reveals novel neuronal architecture in mouse postnatal cortex

We first validated our Large area Spatial Transcriptomic (LaST) pipeline by mapping expression of known cortical layer neuron markers for layers 2-4 (L2-4, *Cux2*), L4 (*Rorb*), L5-6 (*Ctip2*) and L6 (*Foxp2*)(*1*) (Fig 1B,C) in the mouse brain. We performed automated six-color imaging of 10 anterior posterior sections across the entire P14 cortex (Fig 1E), totaling 2,216 fields of view and ~300,000 images collected under 25 hours of continuous acquisition. We used machine learning based texture analysis combined with NeuN IHC (neuronal nuclei and cytoplasm) and DAPI staining to identify single neuron soma on max-z projection images, while also ruling out doublets/triplets with machine-learning (Sup Fig 1, Extended Methods). This approach validated that our technique accurately identified known layer-specific single-cell and layer-specific neuronal gene expression patterns (Fig 1C) with robust reproducibility (Sup Fig 2).

Having generated a quantitative map of known cortical neuron gene expression, we tested whether we could identify and/or discover neuronal subtypes solely from smFISH data in an unbiased manner. We analysed transcript counts from 46,888 neurons across select broad cortical areas using a modified dimensionality reduction and hierarchical clustering approach (Extended Methods, Sup Table 1). Our analysis identified 10 distinct major clusters based on the expression of the four markers above (Fig 1D) and could robustly detect low expression differences across neurons (Sup Fig 3,4,5). These clusters primarily represented layer-enriched neuron populations present across multiple cortical areas (Fig 1C,D,G;, e.g. *Ctip2*^*high*^*Foxp*2^*high*^ cluster #9 in L6). In addition we captured heterogeneity within cortical layers, exemplified by clusters of *Cux2*^*high*^*Rorb*^*low*^ (#1,3-5) and *Cux2*^*high*^*Rorb*^*high*^ (#8) within L2-3. We could also accurately distinguish area-specific differences such as *Rorb*^*high*^*Cux2*^*high*^ neurons (#6 and 8) enriched in L4 of sensory areas and *Cux2*^*mid*^*Ctip2*^*mid*^ neurons (#2, likely an interneuron subtype, Sup Fig 6, (*14*)) in L1-3 and L5 of somatosensory, motor and auditory cortex (full area maps shown in Sup Fig 7). Finally, we found a novel population of *Rorb*^*high*^*Cux*^*mid*^*Ctip2*^*low*^ neurons (#7) restricted to L5 of anterior somatosensory, barrel and visual cortex at P14 and P56 (Fig 1H, I; Sup Fig 6). To confirm the expression of this putative subtype, we examined available cortical single cell transcriptome data (*15*). Consistently, our survey showed that *Rorb* and *Ctip2* expression segregate amongst molecular subtypes of L5 neurons in the adult visual cortex (Sup Fig 6). Taken together, these results demonstrate that LaST map can be used to validate regional qualitative and quantitative predictions about gene expression in mammalian brain and that it can serve as a tool to sensitively identify novel/diversified cell populations *in situ.*

### Laminar gene expression patterns are characteristic of cortical grey matter astrocytes

It is generally accepted that astrocytes in the superficial subpia (L1) and deep L6 show white matter astrocyte characteristics (*16*). We next tested whether gray matter astrocytes in L2-5 are molecularly homogeneous, or alternatively, segment into multiple distinct layers? First, to generate candidate gene lists, we manually dissected the upper (L2-4) and deep (L5-6) layers of the somatosensory cortex from P14 astrocyte reporter *Aldh1L1-GFP* transgenic mice (*17*) and performed RNASeq profiling of FACS-purified astrocytes versus whole cortex (Fig 2A). The expression pattern of whole cortical tissue showed that the microdissection accurately captured upper and deep layers. Astrocyte markers were highly enriched in FACS-purified GFP+ cells over GFP-negative cells and whole cortex tissue as expected; moreover, most neuronal layer markers were not distinctly expressed amongst cortical astrocytes (Sup Fig 8).

**Figure 2:**
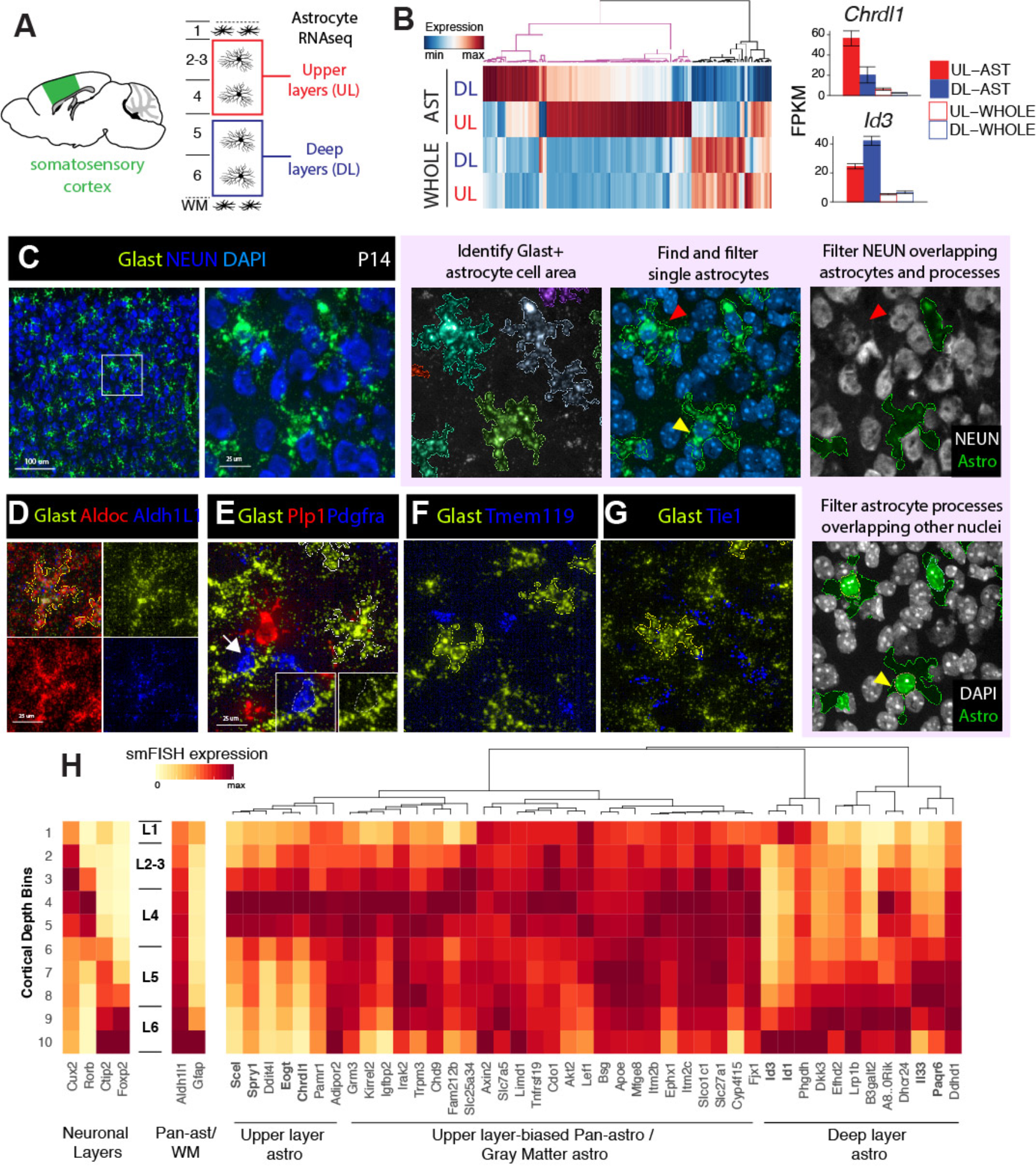
Novel layer expression differences amongst cortical gray matter astrocytes revealed via RNAseq and LaSTmap. **A)** Diagram showing somatosensory cortical areas and layers used for laminar gray matter astrocyte RNAseq expression profiling. **B)** Novel molecular heterogeneity of layer astrocytes identified by RNAseq. 159 differentially expressed genes between upper and deep layer gray matter astrocytes (FDR<0.05 and expression threshold of FPKM>5) were detected (n=3 biological replicates). **C)** Identification of astrocyte gene expression with *Glast (Slc1a3*) smFISH. The astrocyte cell area is segmented from the *Glast* signal and nuclei are segmented from DAPI. Single astrocytes are selected using morphological and intensity filters, overlapping neurons and non-astrocyte nuclei are excluded. **D-G)** *Glast* is a specific marker of astrocytes. Solid outlines indicate the cell areas of identified single astrocytes. (D) *Aldh1l1* expression is too low for clear texture analysis while *Aldoc* expression is too high for astrocyte-specific segmentation. (E-G) *Glast* smFISH with other cell type makers. OPCs (dashed outline) express much lower levels Glast than astrocytes. **H)** Screening candidate layer astrocyte genes with LaSTmap identifies laminar expression patterns *in situ*. Tile heatmaps show average single cell gene expression binned across cortical depth in the P14 somatosensory cortex (n =2 pooled biological replicates across multiple tissue sections). Expression patterns of laminar neuron markers (left), pan-astrocytic marker *Aldh1L1* and white matter astrocyte marker *Gfap* (middle), and candidate layer astrocyte markers from RNAseq profiling (right). Upper and deep layer genes with astrocyte-specific expression are marked in bold. Scalebars: (C, left panel) 100 μm, (all other panels) 25 μm.

This analysis identified 159 candidate genes with a differing expression between the upper and deep layer astrocytes (Fig 2B). While most genes showed moderate-layer enrichment, many also showed layer and astrocyte-enriched expression in published cortical transcriptome datasets (Sup Fig 9). We next validated layer astrocyte gene expression *in situ*. To distinguish astrocyte-specific gene expression by smFISH, we examined markers for astrocytes (*Aldh1l1, AldoC, Glast*), neurons (*Syt1*, NEUN), oligodendrocytes (*Pdgfra, Plp*), microglia (*Tmem119*) and endothelia (*Tie1*) (*17*). As shown in Fig 2C-G, while *Glast* mRNA showed low expression in *Pdgfra+* oligodendrocyte precursors, high *Glast* levels exclusively marked astrocytes. *Glast* mRNA (but not *AldoC* or *Aldh1L1*) filled astrocyte soma and main processes, permitting clear delineation of astrocyte cell borders by image texture analysis.

To avoid double counting, we selected non-overlapping single astrocytes by filtering cells based on size, DAPI (single-nuclei) and *Glast* intensity (Extended Methods, Sup Fig 10 and 11). In addition, we filtered astrocytes and their processes that overlap with neuronal soma and non-astrocyte nuclei in z-projection images (Fig 2C, Extended Methods). These measures ensured our smFISH approach counted single cell astrocyte-specific gene expression *in situ* in the context of a large tissue area screen.

To validate candidate layer-associated astrocyte genes in situ, we first analyzed the P14 somatosensory barrel cortex. We selected the top 46 genes from RNAseq that showed the highest differential enrichment between cortical layers and/or astrocyte specificity (Fig 2H, Sup Table 2). Of these, we confirmed layer enriched expression patterns or rejected genes that showed subtle variations based on quantitative findings *in situ*. For example,% (22/46) showed an expression pattern biased −but not restricted to upper layers (e.g., *Tnfrsf19, Mfge8*), and % (7/46) showed enrichment in all gray matter astrocytes across L2-6 (e.g., *Igfbp2, Kirrel2*). Another third (%, 6/46) showed expression in astrocytes + neurons and were excluded (e.g., *Ddit4l, B3galt2,* Sup Fig 12. In contrast, eight genes clearly distinguished upper versus deep layer astrocytes in the somatosensory cortex (Fig 2H). As shown (Fig 3A-C, Sup Fig 13), expression of the BMP antagonist Chordin-like 1 (*Chrdl1*) (*18*) was expressed in L2-4 astrocytes while Interleukin-33 (*Il-33*) (*19*) was enriched in L5-6. In distinction, Sciellin (*Scel*), a component of epithelial cornified envelopes (*20*), was expressed in mid-cortical (~L4) layers and transcriptional repressors *Id1* and *Id3* occupied deep (L5-6) and marginal (L1) layers (*21*). These findings show that subsets of cortical gray matter astrocytes show laminar spatial gene expression heterogeneity.

**Figure 3:**
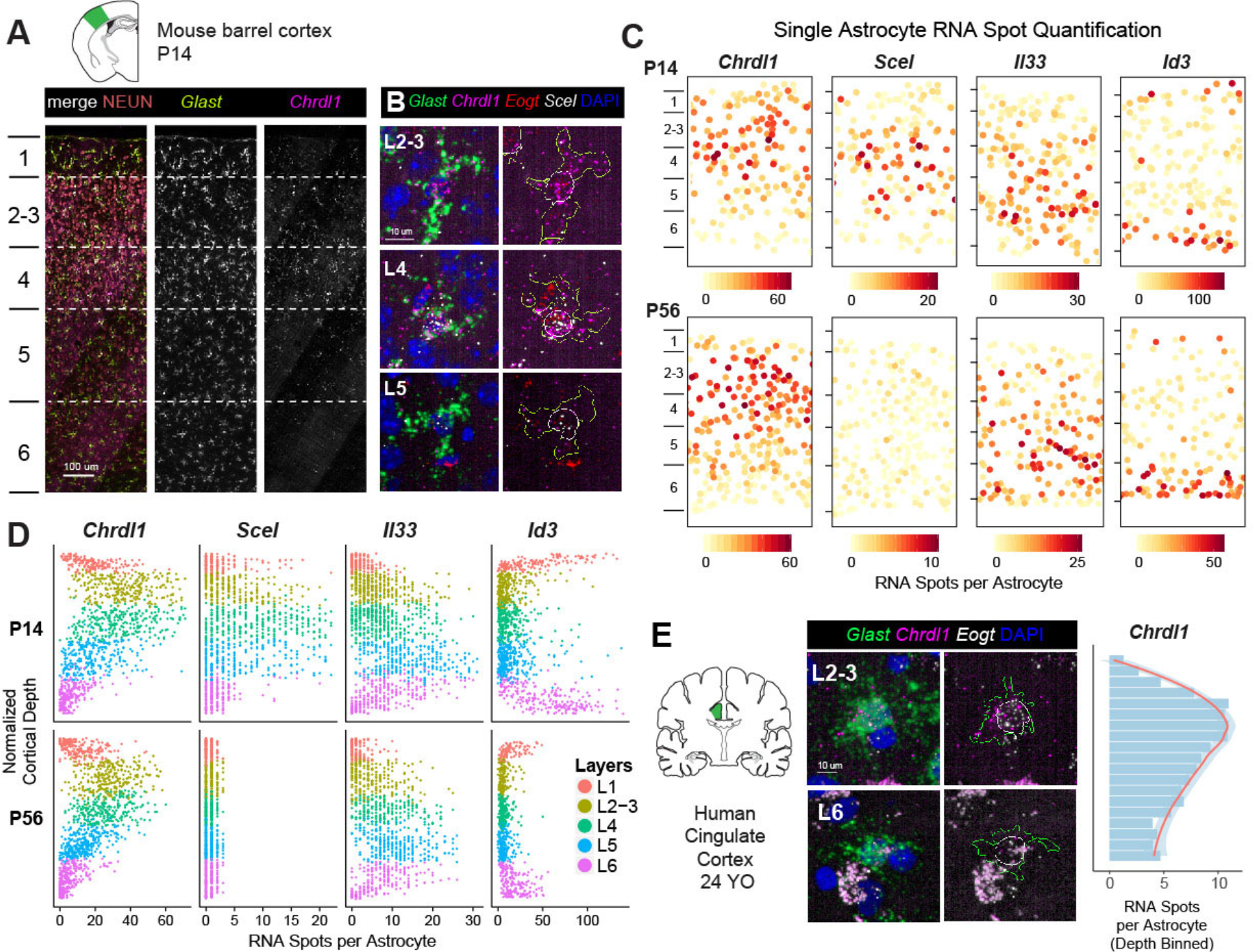
Astrocytes show broad expression gradients across cortical depth and diverge from neuronal layers. **A-B)** Upper layer astrocyte enrichment of *Chrdl1, Scel and Eogt.* Images show the depth of the mouse barrel cortex at P14 (A) and close-ups of astrocytes from different layers (B). Astrocyte cell areas are marked with yellow outlines and nuclei are marked with white outlines. **C)** Astrocytes are organized into superficial, mid and deep layers across the cortical gray matter. Single astrocyte expression maps in the barrel cortex at P14 (top panels) and P56 (bottom panels). Astrocytes are plotted as solid circles and colored quantitively for RNA spot counts per gene per cell. **D)** Single astrocyte quantification of layer astrocyte markers across cortical depth in the barrel cortex at P14 and P56 (n=3 pooled tissue sections from one replicate per timepoint). **E)** Upper layer astrocyte enrichment of *Chrdl1* expression in the adult human cingulate cortex. Quantification of depth binned average single astrocyte expression shown on the right. Scalebars: (A) 100 μm, (B, E) 10 μm.

### Astrocyte layers diverge from neuronal laminae

To determine whether glial layers matched excitatory neurons we directly quantified the spatial expression patterns of the 8 top layer astrocyte markers against layer neuron markers in P14 somatosensory barrel cortex *in situ*. We found that astrocyte layer genes generally fell into three spatial bins comprising superficial, mid and deep laminae. As shown (Fig 3A-D, Sup Fig 13, *Chrld1, Scel, Eogt, Spry1, Paqr6* and *Il33* were expressed in patterns that spanned multiple neuronal layers. Astrocyte *Chrdl1* expression peaked in L2-4, with low levels in L1 and L5. *Scel* expression was highest in L4-5, with an upper boundary in mid L2-3. *Il33* expression peaked in L5, with low levels in L4 and L6. Such patterns were not artifacts of astrocyte size or *Glast* expression level differences across layers (Sup Fig 14). Moreover, analysis of the P56 mouse brain indicated persistence of these patterns except the mid layer *Scel* expression into adulthood (Fig 3D). Indeed, we found that the upper layer astrocyte *Chrdl1* expression was conserved in the adult human brain (Fig 3E, Sup Fig 15).

While we found certain similar features between astrocytes in deep cortical layers and white matter (WM), most cortical astrocyte layer genes were specific to gray matter. L6 astrocytes expressed high levels of *Gfap*, *Id1* and *Id3*, similar to astrocytes in subcortical WM and L1-subpia (Sup Fig 13) (*7*). Conversely, *Il33* was enriched in L5-6 gray matter astrocytes but absent from WM, showing that deep layer astrocyte identity is distinct from white matter. Together, these findings indicate that astrocytes are organized into multiple layers across the cortical gray matter and that several glial laminae are dissimilar to classic neuronal layers (Fig 5E). Moreover, such patterns are evident from P14, persist until adulthood and are conserved in the human cortex.

**Figure 4:**
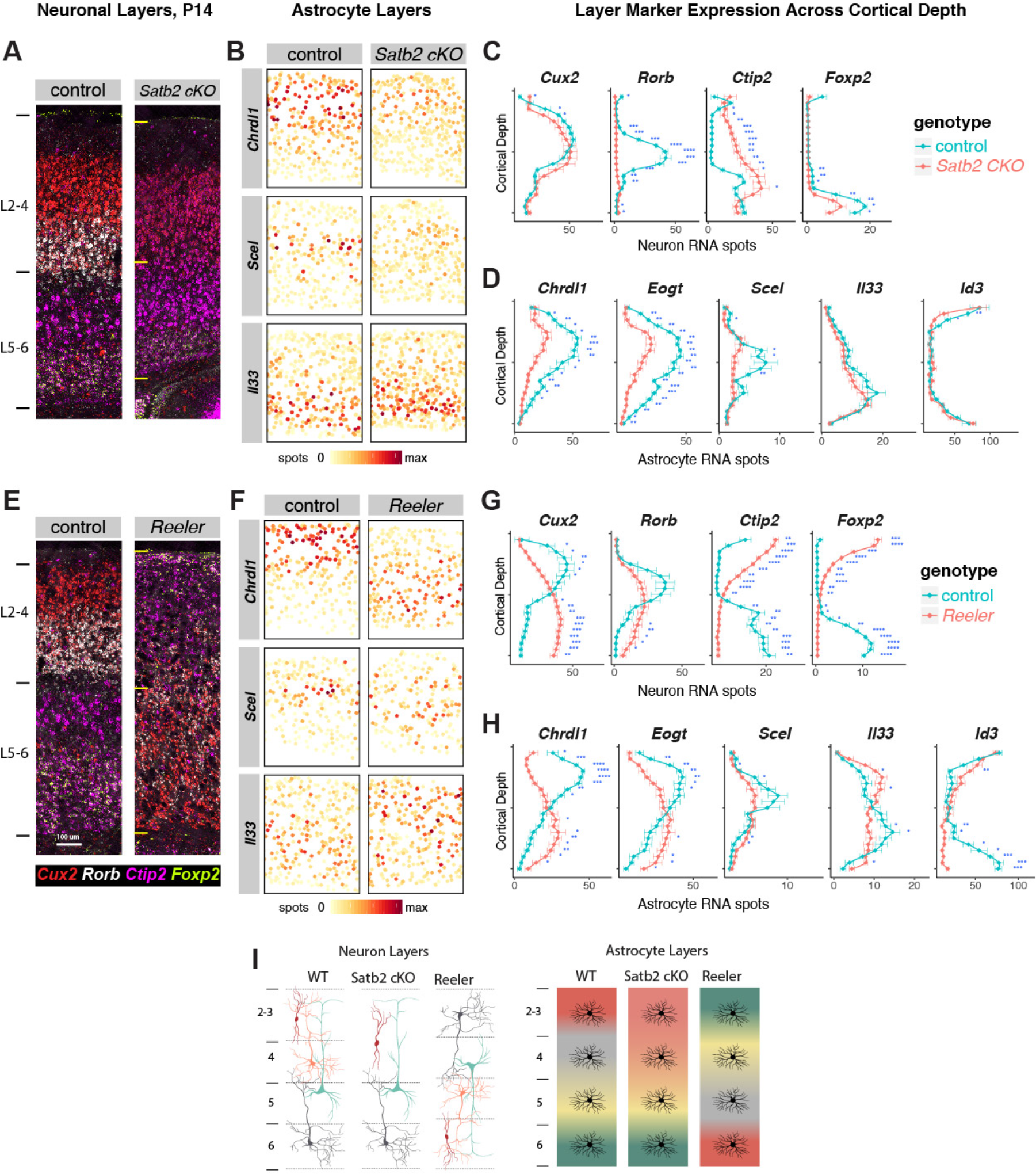
Evidence that post-mitotic neuronal cues establish astrocyte layer identities. **A-D)** *Satb2* cKO mice show defects in upper layer neuron and astrocyte identity. **A)** AImages showing aberrant upper neuronal layers in the *Satb2* cKO barrel cortex at P14. **B)** Single astrocyte maps comparing layer astrocyte marker gene expression in *Satb2* cKO to littermate controls. **C)** Quantification of cortical depth binned neuronal layer marker expression in cKO vs control (n=3 pooled biological replicates per genotype). *Satb2* cKO shows loss of L4 neuron gene expression and some ectopic deep layer marker expression in L2-3. **D)** Quantification of cortical depth binned astrocyte layer marker expression in cKO vs control (n=3). *Satb2* cKO shows complete loss of mid-layer astrocyte Scel expression and reduction in upper layer astrocyte markers *Chrdl1* and *Eogt*. **E-H)** *Reeler* mice show inversion of neuronal and astrocyte layers. **E)** Images showing neuronal layer inversion in the *Reln*-/ barrel cortex at P14. **F)** Single astrocyte maps comparing layer astrocyte marker gene expression in *Reln-/-* to littermate controls. **G)** Quantification of cortical depth binned neuronal layer marker expression in KO vs control (n=3 pooled biological replicates per genotype). *Reln* shows inversion of superficial and deep neuronal layers with L4 neurons forming aberrant clusters across the cortex. **H)** Quantification of cortical depth binned astrocyte layer marker expression in cKO vs control (n=3). *Reln* shows inversion of superficial and deep astrocyte layers with mid layer astrocytes are scattered across the cortex. **I** Diagrams depicting layer neuron and astrocyte changes in *Satb2* cKO and *Reln-/-* mice. Scalebars: 100 μm. All data represent mean ± s.d. *P < 0.05,**P < 0.01, ***P < 0.001.

**Figure 5:**
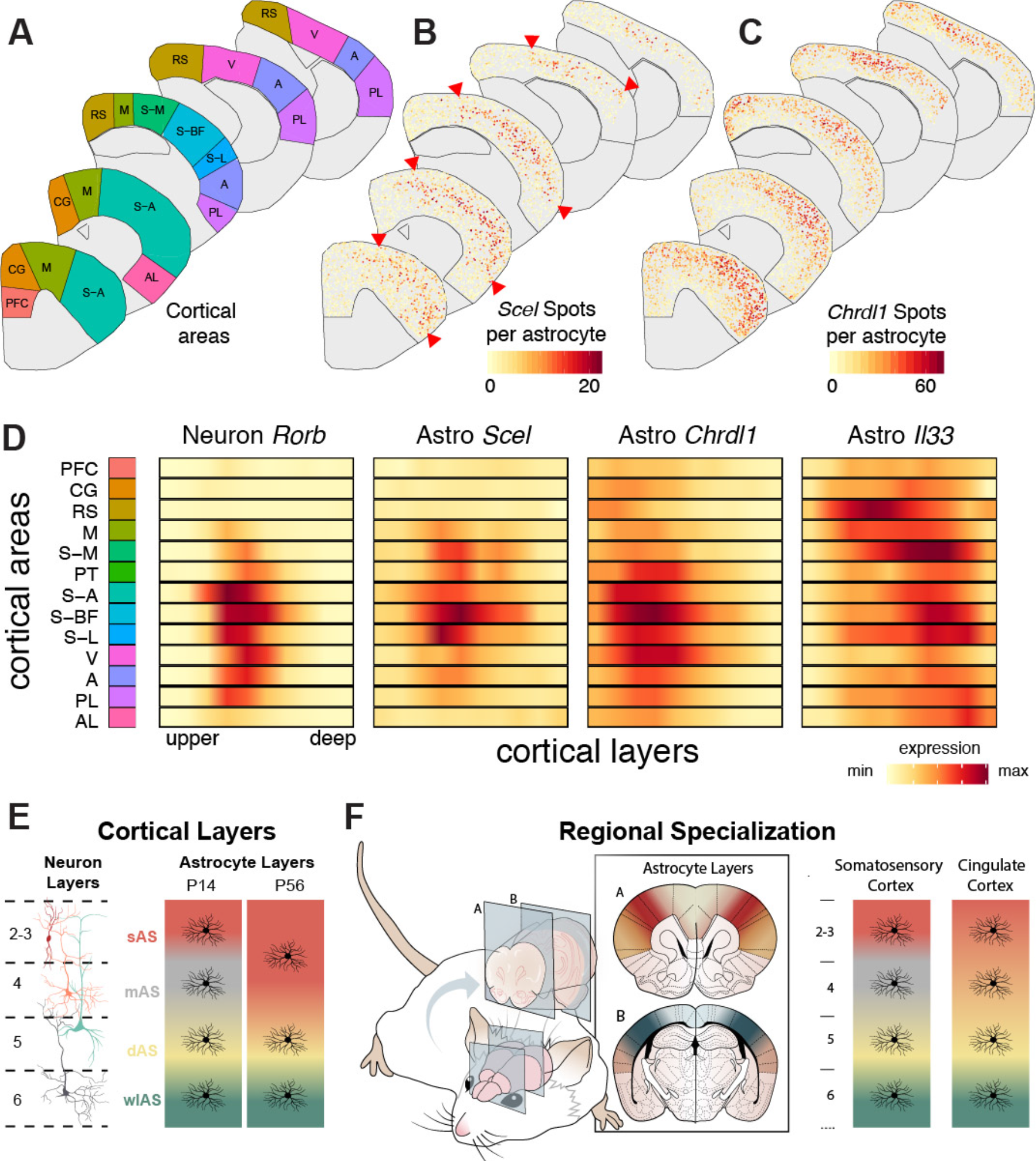
Astrocyte arealization across the cortex. **A-C)** Single cell mapping of astrocyte gene expression across cortical areas. Maps show the (A) P14 cortical areas used in the analysis, (B) single astrocyte expression of *Scel* and (C) *Chrdl1* across the dorsoventral and rostrocaudal extent of the cortex. Astrocytes are plotted as solid circles and colored quantitively for RNA spot counts per gene per cell. Arrowheads indicate the restriction of *Scel* expression to sensory areas. Abbreviations as in Fig 1. **D)** Astrocyte layers are distinct across cortical areas. Smoothened tile plots showing the quantification of neuronal *Rorb* expression and astrocyte expression of *Scel, Chrdl* and *Il33* across the cortical areas and layers (n=10 tissue sections from one biological replicate). **E)** Diagrams showing the divergent layer heterogeneity of astrocytes. Our study identifies superficial (sAS), mid (mAS) and deep (dAS) layer astrocyte subtypes through cortical gray matter in the postnatal cortex and confirms white matter like (wlAS) properties of L6 astrocytes. sAS, dAS and wlAS layer heterogeneity are maintained into adulthood. **F)** 3D model showing astrocyte area and layer heterogeneity. Astrocyte layering is regionally specialized across the dorsoventral and rostrocaudal extent of the cortex. sAS and mAS subtypes are enriched in sensory areas while dAS subtypes are enriched in medial areas.

### Neuronal specification directly or indirectly determines astrocyte laminar gene expression

What determines astrocyte layer gene expression? During cortical development, neuronal layers are established along a radial glial scaffold prior to astrogenesis (P0-7 (*22*)). To investigate whether neuronal factors regulate astrocyte layer organization, we first examined mice that lack the chromatin remodeling factor *Satb2*, which is required to specify superficial post-mitotic callosal neuron identity (*23*). *Satb2* conditional knockout mice (*Satb2* cKO*, Emx1-cre x Satb2 floxed/floxed*) are deficient in L2-4 neuron gene expression, lack callosal axon projections and ectopically express markers of L5-6 neurons in superficial cortical layers (*24, 25*).

We verified that at P14, L4 neurons (*Rorb, Pamr1*) are absent in *Satb2* cKO while L2-3 neurons show some ectopic deep layer gene expression (*Ctip2*) (Fig 4A,C, Sup Fig 16). As a consequence, we found complete loss of *Scel* expression in astrocytes, while upper layer astrocyte genes *Chrdl1* and *Eogt* were significantly reduced (Fig 4B,D). Deep layer astrocyte genes (*Il33, Id1, Id3*) were unaffected. These findings indicate that post-mitotic L4-specific neuron identity is necessary for the generation of superficial layer astrocyte identity (Fig 4I). Next to assess deep layer astrocyte identity in relation to neurons, we used *Reeler (Reln*) mice, deficient for secreted Reelin, that show an inversion of radial glial polarity and neuronal layers (*26*). In *Reln*, excitatory neurons are specified correctly but fail to migrate to their proper layers, yielding an “inside-out” cortex before birth (*27*). As shown at P14 (Fig 4E-G), the expression pattern of superficial (*Cux2*) and deep laminar markers (*Ctip2, Foxp2*) were inverted while L4 neurons (*Rorb*) formed aberrant clusters across cortical depth, as previously shown (*28*). As a consequence, we found that superficial and deep astrocyte layers were also inverted (Fig 4F-H). Consistent with the L4 neuron phenotype, Scel+ astrocytes were aberrantly located across cortical depth. These findings suggest roles for cortical neuronal cues in regulation of astrocyte layer development (Fig 4I).

### Astrocyte layer characteristics vary between cortical areas

Are astrocyte layers distinct or similar between functional cortical areas? To investigate astrocyte arealization, we quantified patterns of top layer astrocyte markers across the P14 cortex. As shown (Fig. 5), astrocyte layer gene expression varied significantly across dorsoventral and rostrocaudal cortex. *Scel* expression showed enrichment in L4-5 astrocytes in the somatosensory areas but was absent from the medial motor and caudal visual cortex (Fig 5A,B), a pattern that correlates closely with L4 neurons (Fig 5D). In a different pattern, *Chrdl1* showed dorsoventral enrichment in L1-4 astrocytes throughout the cortex (Fig 5C). In contrast, deep layer astrocyte marker expression was enriched in medial and lateral areas across the dorsoventral axis (*Il33, Id3*, Fig 5D, Sup Fig 17, 18). Thus, astrocytes show both laminar and area heterogeneity across the cortex. Collectively, our observations on the regional heterogeneity of cortical astrocytes and their induction by neuronal signals show that glial laminae are a fundamental biomarker characteristic of cortical layer and area architecture (Fig 5E,F).

## Discussion

International projects such as the human cell atlas and BRAIN initiative have generated large repositories of gene expression data, and a logical next step is to map back such expression patterns *in situ* within developing tissues and/or mature organs. However, spatial transcriptomic analysis in human tissue presents a particular scaling challenge. LaSTmap provides a pipeline to map quantitative gene expression in single cells across large mouse and human tissue areas. It uses commercially available probes and does not require special tissue preparation or bespoke imaging equipment. LaSTmap showed reproducibility across multiple tissue samples/sections and should be adaptable for a range of other tissues to obtain regional/qualitative and cellular/quantitative information from ISH reintegrated into 2D tissue maps. Future elaboration with tissue warp integration would enable *in silico* 3D map construction (*29*).

Our findings indicate LaSTmap is a robust technique to validate single cell gene expression *in situ* as well as discovery of combinatorial patterns indicative of cell diversity. In particular, our finings indicate a novel Rorb/Ctip2 L5 neuronal sub-type in visual cortex, as well as molecular diversity of cortical gray matter astrocytes. Indeed, a surprising biological finding of our study is that cortical gray matter astrocytes are organized into lamina that are distinct from classic layers of excitatory neurons. Our study identifies the segmentation within astrocytes across L2-5, indicating specialized gray matter sub-types. In our study, the quantification of astrocyte layer genes uniquely showed peaks and troughs of expression across the superficial-to-deep cortex, as well as differences across functionally distinct cortical areas. As prior studies show that region restricted astrocyte mRNA and protein expression are predictors of functions tailored to support of local neural circuits (*8*, *30*, *31*), astrocyte laminar genes indicate potential additional localized functions (*32*).

Analysis of cortical lamination mutants suggested that neuronal cues are required, directly or indirectly, to establish astrocyte laminae, particularly in L4 where neurons and mid-layer astrocyte layers do match most closely. However, we cannot rule out that a cortical ‘pre-pattern’ instructs astrocyte precursors to respond to neuronal signals or other mediator cells involved in astrocyte specification. Indeed, astrocyte-encoded gradient patterns are typical of cells that have undergone polarization in response to a morphogen, and upstream regulation of this process is a subject for further study. Together, our findings indicate the diversification of astrocytes in layer patterns that diverge from excitatory neurons, indicating a composite “neuroglial” cortical architecture with higher order complexity.

## Author Contributions

O.A.B. and D.H.R. conceived the study. O.A.B. planned the experiments. O.A.B., T.B. and S.H. performed histology and imaging experiments. O.A.B. performed the image analysis. K.P. and J.S. contributed to the histology and imaging pipeline. D.P. and O.A.B. analyzed neuron gene expression data to identify subtypes. A.B. wrote the slideSegmenter software. A.M.H.Y. and M.F.P. provided human tissue. O.A.B. and L.B.H. performed layer astrocyte purification. O.A.B. and R.K. analyzed RNASeq data. K.S. and S.M. supported mouse work and genotyping. O.A.B. and D.H.R. wrote the manuscript with feedback from all authors.

## Acknowledgments

We are grateful to Ben Barres and Sarah Teichmann for helpful discussions and comments. We are grateful to Barry Lynch and Xiao-Jun Ma for advice on RNAScope, as well as Richard Sawkins and James Hutt for technical support on imaging. We are also grateful to Eric Olson (SUNY Upstate University) for providing *Reeler* mice and Ralph Marcucio (UCSF) for providing the *Satb2*-*flox* mice.

## Fellowships

The authors were supported by the Life Sciences Research Fellowship and the Howard Hughes Medical Institute (to O.A.B.), the Wellcome Trust (to T.B.), NIHR Academic Clinical Fellowship and a Wellcome Trust PhD for Clinicians Fellowship (to A.M.H.Y.)

## Supplementary Methods

### Mice

All mouse strains were maintained at the University of California, San Francisco (UCSF) specific pathogen-free animal facility and all animal protocols were approved by and in accordance with the guidelines established by the Institutional Animal Care and Use Committee and Laboratory Animal Resource Center. The day of birth was designated as Postnatal day 0 (P0).

Unless otherwise started, wild-type Swiss Webster mice were used for histology and spatial transcriptomics. *Aldh1l1-GFP* transgenic mice were generated by the GENSAT project (*33*). *Emx1-cre* mice were obtained from The Jackson Laboratory (JAX #005628) (*34*). Satb2-flox mice were a gift from Dr. Ralph Marcucio at UCSF (*25*). Conditional knockouts were generated by breeding *Emx1-cre/+; Satb2-flox/+* with *Satb2-flox/flox* mice to obtain mutants (*cre/+; flox/flox*) and littermate controls (*cre/+;flox/+*). *Reeler* (*Reln/Reln*) mice were a gift from Dr. Eric Olson at SUNY Upstate University (B6C3Fe a/a-Reln; JAX). Reln/+ mice were used as littermate controls. All mice were maintained on a mixed background.

### Mouse tissue preparation

Mice were transcardially perfused at P14 or P56 with ice-cold phosphate buffer saline (PBS) and 4% paraformaldehyde (PFA) in 1X PBS. Brains were dissected and post-fixed in 4% PFA for 24 hours at 4°C. Post-fixed brains were cryo-protected in 30% sucrose for 48 hr at 4°C and embedded in optimal cutting temperature compound (Tissue-Tek). Cryosections (16 microns) were collected on superfrost slides (VWR) using a cryostat (CM3050S, Leica) and stored at −80°C.

### Human tissue

Human brain tissue was obtained with informed consent under protocol 16/LO/2168 approved by the NHS Health Research Authority at the Addenbrookes Hospital. Adult brain tissue biopsies were taken from the site of neurosurgery resection for the original clinical indication. For the purposes on this study samples were taken from peri contusional areas in traumatic brain injury (frontal cortex), lobectomy in epilepsy surgery (temporal cortex) and peri-tumoural tissue (temporal cortex).

Tissue specimens were collected in Addenbrookes Hospital and transferred to a CL2 facility where it was processed. Tissue was dissected and fixed in 4% PFA for 48-72 hours. Once fixed, samples were placed in 20% Sucrose for cryoprotection for 24-48 hours and mounted in OCT, stored at −80C.

Additional human brain tissue was collected in a de-identified manner with previous patient consent in strict observance of the legal and institutional ethical regulations of the University of California, San Francisco (UCSF) Committee on Human Research. Protocols were approved by the Human Gamete, Embryo and Stem Cell Research Committee (Institutional Review Board) at UCSF. For this study, one post mortem sample was taken from the superior frontal gyrus and processed as above. Human brain blocks were cryosectioned to 16 microns.

### smFISH assay design and probes

Mouse and human tissue smFISH was performed using the RNAScope LS Multiplex Assay (Advanced Cell Diagnostics, ACD) (*35*). In this assay, the smFISH signal-to-noise ratio (SNR) is amplified by branched DNA complexes formed on target transcripts and tyramide signal amplification (TSA)-based labeling. Target RNAs are initially hybridized to a series of single-stranded DNA “z-probes”. Each z-probe is composed of (**1**) a 18-25 nucleotide region complementary to the target RNA, (**2**) a spacer sequence, and (**3**) a 14 nucleotide tail region. These probes are tagged by branched DNA-amplification trees: pairs of z-probes are hybridized to oligo-preamplifiers, across their bridged tail sequences, which are then tagged by 20 oligo-amplifiers (Fig 1 A). Each oligo-amplifier is labeled with 20 Horse Radish Peroxidase (HRP) enzyme molecules. In general, a 1 kilobase region on the target transcript is hybridized by 20 z-probe pairs in tandem, which can yield up to 8000 HRP labels per each target. The fluorescent smFISH signal is consequently generated by the addition of tyramide-conjugated fluorophores. Tyramide is enzymatically converted into a highly oxidized intermediate by HRP that covalently binds to the proteins at or near the HRP label, depositing a large number of fluorophores for probe detection(*36*). The combination of branched DNA and TSA-amplification significantly boosts the sensitivity and SNR of the RNAScope assay, allowing fast confocal imaging of large tissue areas with short exposure times (see imaging).

#### Multiplexed detection

To achieve 4-plex transcript detection with RNAScope, target z-probes are assigned to one of four different channels (C1-C4) that contain distinct tail-sequences. Tissue samples are hybridized to the mixture of C1-C4 probes, followed by generation of channel-specific amplification trees. Finally, probes are sequentially developed with TSA through incubation cycles of channel-specific HRP labels, tyramide-conjugated fluorescent dyes and chemical enzymatic quenchers.

#### Probe information

All of the RNAscope probes used in this study and relevant information including target sequences are listed in Supplementary Table 3. Further information is readily available from the vendor (https://acdbio.com/catalog-probes). To assess the background signal from the assay, target probes against the bacterial *DapB* mRNA were used as negative controls. Target probes against mouse and human housekeeping genes were used as positive controls. In the mouse cortical astrocyte expression screen, the *Glast* probe was always assigned to C4 and multiplexed with other probes in C1-C3 channels. With human tissue, the *Glast* probe was assigned to C3 and multiplexed with C1-C2.

### Automated smFISH and IHC

All histology on mouse and human brain cryosections was automated on a Leica BOND RX robotic stainer after manual baking and dehydration. Tissue sections were first processed for 3 or 4 gene smFISH using the RNAScope LS Multiplex Assay (ACD). After smFISH, antibody staining was performed using TSA and slides were manually coverslipped for imaging.

#### i. Baking and dehydration

Tissue cryosections were removed from −80°C and thawed at RT for 15 min. Samples were then baked at 65°C for 45 min in vertical position on a slide holder (Tissue-Tek) in an oven. After baking, samples were dehydrated in a series of 50%, 70%, 100% and 100% ethanol (5 min each) in staining dishes (Tissue-Tek) and air-dried for 10 min before automated RNAScope.

#### ii. Automated histology design and setup

For use on the Leica BOND RX, all histology consumables were transferred to barcoded reagent containers. Staining protocols that list the order and durations of reagent incubations and washes were created on the BOND controller software. Slides were assigned unique barcode labels coupled to staining protocols and placed onto temperature-controlled pads on the instrument. Flow through chambers were assembled across the whole slides using plastic coverplates. During staining, a liquid volume of 150 µL was dispensed to each slide on every step using automated liquid handling. Reagents were flushed at least once before main incubations to ensure uniform coverage of the slide. Between reagent incubations, multiple short washes were performed. All incubations were performed at room temperature unless indicated otherwise. For the 4-plex RNAScope smFISH protocol, a maximum number of 20 slides could be processed against 10 different probe mixtures in a single run (e.g. 40 different genes screened across two sets of biological replicates). The combined multiplexed RNAScope smFISH and IHC protocol for 20 slides ran overnight on the Leica BOND RX lasting ~17 hours.

#### iii. Automated RNAScope smFISH

The RNAScope LS Multiplex assay (ACD) was performed largely according to the instructions from the vendor and modifications are noted below. Full details of the protocols 4-plex smFISH and the consumables used in this study are provided in Supplementary Table 4.

##### 4-plex probe hybridization

To perform 4-plex RNAScope on mouse brain cryosections, samples were initially permeabilized with heat and protease treatment to improve probe penetration and hybridization. For heat treatment, P14 and P56 samples were incubated in BOND ER2 buffer (pH 9.0, Leica) at 95°C for 2 and 5 min, respectively. For protease treatment, P14 and P56 samples were incubated in ACD protease reagent at 42°C for 10 and 15 min, respectively. Prior to probe hybridization, samples were incubated in hydrogen peroxide for 10 min to inactivate endogenous peroxidases and ACD protease. Following pre-treatment, samples were incubated in target z-probe mixtures (C1-C4) for 2 h at 42°C. The C2-C4 probes are provided at 50X concentration by ACD and were diluted 1:50 in C1 probes. In exception, the C4 probe for the high-expressing *Glast* mRNA was used at 1:100 in the astrocyte screen for reagent conservation.

##### smFISH signal amplification

After probe hybridization, branched DNA amplification trees were generated through sequential incubations in AMP1, AMP2 and AMP3 reagents for 15-30 min each at 42°C with LS Rinse buffer high stringency washes between incubations. Following amplification, probe channels were detected sequentially via HRP-TSA labeling. To develop the C1-C3 probe signals, samples were incubated in channel-specific HRP reagents for 15 min at 42°C, TSA fluorophores for 30 min and HRP blocking reagent for 15 min at 42°C. The probes in C1, C2 and C3 channels were labeled using Opal 520, 570 and 650 fluorophores (Perkin Elmer, diluted 1:2500) respectively. Finally, to develop the C4 probe, the Atto425 fluorophore was used for 6 color imaging on the Operetta system. The C4 probe complexes were first incubated with TSA-biotin (Perkin Elmer, 1:500) for 30 min, followed by streptavidin-conjugated Atto425 (Sigma, 1:400) for 30 min. Multiple short washes were performed between incubations throughout the protocol using the BOND Wash buffer (Leica) and deionized water (full protocol listed on Supplementary Table 4).

##### 3-plex smFISH

To perform 3-plex RNAScope on human brain cryosections, samples were heat-treated for 10 min and incubated in protease for 15 min. Probe hybridization and branched DNA amplification were performed as described above. To develop C1-C3 probes, Opal fluorophores (520, 570 and 650) were used at a lower dilution (1:300) due to higher autofluorescence on postnatal human brain sections. To distinguish RNA spots from lipofuscin autofluorescence, spots that appear identical across Opal 520 and 570 channels were filtered out.

##### iv. Automated immunohistochemistry

RNAScope smFISH was directly followed by antibody staining for the neuronal marker NEUN on the BOND RX system. To improve antibody staining after IHC and perform 6-color imaging on the Operetta, the NEUN signal was amplified using TSA-biotin and the Alexa 700 fluorophore. Samples were first blocked in antibody blocking solution (Perkin Elmer) for 20 minutes. To block any available TSA-biotin sites from the smFISH assay, samples were incubated in 0.2% Avidin (Sigma) for 20 min and 0.05% Biotin (Sigma) for 30 min. After the avidin-biotin block, samples were incubated in chicken anti-NEUN antibody (Milipore) diluted 1:500 in blocking solution for 1 hr. To develop the antibody signal, samples were incubated in goat anti-chicken HRP (ThermoFisher, 1:500) for 1 hr, TSA-biotin (1:200) for 10 min and streptavidin-conjugated Alexa 700 (Sigma, 1:200) for 30 min. Following antibody staining, samples were incubated in DAPI (Sigma, 0.25 µg/ml) to mark cell nuclei and washed multiple times in deionized water. After final washes, slides were briefly air dried and manually mounted using ~170 µL of Prolong Diamond Antifade (Fisher Scientific) and standard coverslips (24×50 mm; Fisher Scientific). The full IHC protocol is listed under Supplementary Table 4.

### Automated spinning disk confocal imaging

Tissue sections were imaged on an Operetta CLS high-content screening microscope (Perkin Elmer). To perform 6-color smFISH-IHC imaging, this system was equipped with 8 LED light sources, 5X air and 40X water objectives, wide-field and spinning disk confocal imaging modules and narrow band emission filters. The fluorophores, light sources, exposure times and emission filters used for mouse and human tissue imaging experiments are listed in Supplementary Table 5. Image acquisition and analysis were controlled using the Harmony software (Perkin Elmer).

#### i. Tissue identification

To locate whole tissue sections or ROIs for high-resolution imaging, entire slides were initially scanned under low magnification in wide-field mode. Each slide was imaged for nuclear DAPI and NEUN staining if applicable using a 5X NA 0.16 objective (pixel size: 7.2 μm) under 5 minutes. To automatically locate the xy coordinates of tissue sections, a Harmony analysis script was used to detect DAPI+ areas. Whole slide DAPI images were stitched, smoothed with Gaussian blurring and analyzed with a global threshold. The detected DAPI+ areas were size filtered to remove staining artifacts and slightly expanded to ensure complete tissue coverage. The resulting areas were used to automatically set the xy-field positions of the subsequent 40X scan. Alternatively, ROIs for 40X scans were manually selected on low magnification previews. Selected 40X fields were imaged with a 7% overlap.

#### ii. Confocal imaging

The high-resolution smFISH images of tissue sections were acquired on the spinning disk confocal mode using a sCMOS camera and a 40X NA 1.1 automated-water dispensing objective. The field-of-view was 320 x 320 μm and the pixel size was 298 nm. A P14 mouse brain hemisection comprised 200 to 300 fields depending on its anatomical position. Each field was imaged as a z-stack consisting of 20 to 30 planes with a 1 μm step size across each color channel. An IR laser was used to auto focus on the position of the coverslip and the relative z-heights of tissue sections were manually identified by imaging DAPI on sample fields prior to tissue-wide scans. Each z plane was imaged across 4-6 channels depending on the experiment with exposure times for mouse smFISH channels between 60 and 120 ms (Supplementary Table 5). The 40X multi-channel settings and tissue heights were entered into an experimental layout on Harmony and automatically executed after low magnification tissue identification scans.

### Image analysis

To segment single neurons and astrocytes and quantify RNA spots from high-resolution images, analysis scripts were created on Harmony software (Perkin Elmer) using customizable pre-defined function blocks (*italicized below*). The complete single neuron and astrocyte segmentation pipelines are provided in Supplementary Tables 6 and 7. Each 40X field was analyzed separately to optimize processing time of large datasets.

#### i. Quantification of neuronal gene expression *in situ*

##### a. Segmentation of neurons

Maximum intensity z-projection images were calculated across each channel to generate 2D images from z-stacks. NEUN+ neurons were segmented in three steps. **1**) Supervised texture segmentation was performed at a coarse scale to locate NEUN+ areas on images and filter staining artifacts on tissue sections (*find texture regions*). Intensity and size thresholding then identified the NEUN+ neuronal soma (*find image region*). **2**) Neuronal nuclei were segmented within the NEUN+ soma from Gaussian blurred DAPI images using intensity, size and contrast thresholds (*find nuclei*). **3**) The neuronal cytoplasm was segmented around the nuclei within the boundaries of the neuronal soma using NEUN intensity thresholding (*find cytoplasm*).

##### b. Filtering single neurons

Neuron segmentation yielded single neurons as well as doublets/triplets that overlap in z-projection images and neurons that are partially contained in tissue sections (Sup Fig 1B). To automatically distinguish single neurons, morphological (e.g. area, roundness) and intensity (e.g. pixel sum over DAPI and NEUN) properties of segmented cells were measured and used to train a supervised linear classifier (*select population*). For the training set, we manually selected more one hundred single, doublet and partial cells across multiple tissue sections and cortical areas. The resulting classification was validated across the cortex by manual inspection of several fields.

##### c. RNA spot calling in neurons

RNA spots were identified by the detection of local intensity maxima across each smFISH channel in the neuronal soma (*find spots*). Individual spots were identified with an upper radius threshold of 750 nm. The number of RNA spots per single neuron was calculated for each smFISH channel (*calculate properties*). Last, all DAPI+ nuclei were identified across the given field for use in brain region segmentation (see anatomical annotation below).

#### ii. Quantification of astrocyte gene expression *in situ*

##### a. Segmentation of astrocytes

Maximum intensity z-projection images were generated as above. Background illumination profiles of fluorescent channels were mapped to correct uneven illumination (*flatfield correction*). *Glast*+ astrocytes were segmented in three steps: **1**) Supervised texture segmentation was performed at a fine scale to identify *Glast*+ astrocyte cytoplasm and main processes (*find texture regions*). To train texture analysis, over a hundred points were selected inside versus outside *Glast*+ cortical astrocytes across several tissue sections. Astrocyte cell areas were then filtered by size to remove partial cells and holes across astrocyte nuclei, which are weakly labeled by *Glast* smFISH, were filled (*select region*). **2**) Astrocyte nuclei were segmented within *Glast*+ cell areas from Gaussian blurred DAPI images using intensity, size and contrast thresholds (*find nuclei*). To remove false positive non-astrocyte nuclei that overlap with astrocyte processes in z-projections, additional morphology and Glast intensity filters were used (*select population*). **3**) The astrocyte cytoplasm and processes were segmented around the nuclei within the cell areas using *Glast* intensity thresholding (*find cytoplasm*).

##### b. Filtering single astrocytes

To remove overlapping astrocyte doublets and partial cells, cells were filtered based on morphological (e.g. area, roundness) and intensity (e.g. pixel sum over DAPI and *Glast*) properties. As shown in Sup Fig 11, upper layer astrocytes were slightly larger consistent with previous reports (*7*) and showed higher *Glast* levels than those in deep layers. This analysis identified similar numbers of astrocytes across cortical areas in technical replicates (Sup Fig 11).

Given their intimate cell-cell interactions, astrocyte processes occasionally overlapped with neuronal soma and other nuclei in maximum-z projection images. To remove overlapping non-astrocyte nuclei, the DAPI signal in the astrocyte cytoplasm was identified with intensity thresholding and subtracted from the *Glast+* cell area (cytoDAPI filtered cells, Fig 2C). To remove overlapping neurons, NEUN signal was used to segment neurons as above. Astrocytes that significantly overlap with neurons were discarded (>50% overlap between the astrocyte nuclei and neuronal soma). NEUN+ neurons were then subtracted from the cytoDAPI-filtered Glast+ cell area (Fig 2C), resulting in astrocytes filtered against overlapping neurons and nuclei.

##### d. RNA spot calling

RNA spots were quantified across single astrocytes as elaborated for neuron previously. In addition to the filters used above, cells that are high-expression outliers (above 99.5% of spot counts per gene across the brain) were also filtered out. RNA spots in neurons, cytoDAPI-filtered and double-filtered astrocytes were also quantified.

#### iii. Data processing

The analysis was performed on a desktop workstation with two 6-core Intel i7-4930K 3.4 GHz CPUs and 64 GB of RAM. The neuronal dataset shown in Fig 1, consisting of 10 tissue sections and ~300,000 images, was analyzed under 6 hours. The data was exported from Harmony as a single cell matrix showing cell coordinates, morphological and intensity measurements, and RNA spot counts per cell. Brain region segmentation was performed in MATLAB (described below). Data organization and plotting were done in R.

### Anatomical annotation of mouse cortical areas

For mapping neuron and astrocyte subtypes across cortical areas, one P14 mouse brain hemisphere was sectioned along the coronal plane to generate an 8-slide series containing 10 sections each. One slide was used to map the expression of cortical layer neuron markers (Fig 1) and the remaining slides were assayed with layer astrocyte markers (Fig 3 and 5). The cortical areas were annotated using NEUN and DAPI staining as well as layer neuron marker expression patterns as anatomical landmarks (Sup Fig 7). The Paxinos (*P6, plates 9 to 40*) and Allen Mouse Brain ISH Atlases (*P56, sections 38 to 88*) were used as anatomical references. Cortical areas were annotated broadly across the anterior-posterior and dorso-ventral axes, grouping functionally related areas (e.g. the anterior division of the somatosensory cortex contains the mouth and limb areas). The list of cortical area abbreviations and groupings are listed under Sup Table 8.

For screening layer astrocyte gene expression (Fig 2), two P14 mouse brain hemispheres (biological replicates) were sagitally sectioned to generate 18 slides. Each slide contained 4 sections through the somatosensory cortex from each replicate, corresponding to the areas used for RNAseq profiling of layer astrocytes.

For examining cortical layers in *Satb2* cKO and *Reeler* mice (Fig 4), three coronal sections through the somatosensory barrel cortex were collected from littermate control and mutant brains (n=3 biological replicates each). Each slide contained sections from one control and one mutant brain.

### Manual segmentation of brain regions

Brain areas were manually segmented on low magnification (5X) images of DAPI/NEUN stained brain sections. These segmentation masks were overlaid on xy-coordinates of high-magnification (40X) scans to annotate single cells. The offset between 5X and 40X objectives was manually corrected by aligning DAPI+ nuclei (identified across all cells in each field at the end of segmentation pipelines). To stitch images from Harmony, draw and name segmentation masks, align low-high magnification data and perform batch segmentations, the slideSegmenter application was created to work on the MATLAB environment and made publicly available (https://bitbucket.org/alexmatlab/slidesegmenter/).

### Identification of neuronal subtypes from smFISH data

To identify cortical neuron subtypes in an unbiased manner from smFISH data, we adopted the following workflow:

#### i. Filtering and normalization

For downstream analysis, neurons were 1) selected from the 8 broad cortical areas that show the full complement of layers with respect to the 4 genes profiled; and 2) filtered with a minimum cumulative spot-count threshold of 20. Spot counts then had a value of 1 added and were log transformed (log(spot counts +1)).

#### ii. Clustering

Clustering was performed for cells from each region individually with the 4 genes profiled using graph-based clustering implemented by Seurat (FindClusters function, resolution 0.5) (*37*). Briefly, a K-nearest neighbor graph based on Euclidean distance is constructed from the expression values for each cell. Edges between cells were weighted based on shared overlap in neighborhoods determined by Jaccard distance. Cells were iteratively grouped together with the goal of optimizing the density of links inside communities as compared to links between communities.

#### iii. tSNE

For visualization, t-distributed stochastic neighbor embedding (tSNE) coordinates were calculated from the expression values for each cell (independent of the clustering) using perplexity 250 with Seurat (RunTSNE function). tSNE plots were then colored by the cluster assignments derived above, gene expression values, or other features of interest.

#### iv. Hierarchical clustering

The mean expression profiles of each of the Seurat clusters derived from each brain region were taken and hierarchically clustered together based on Euclidean distance using Ward.D2 clustering (hclust(dist, method = “ward.D2”) R function). The resulting dendrogram was then cut at height 1.9 yielding 18 groups (cutree(hc, h = 1.9) R function). These groups were manually annotated to 10 major subtypes based on high expression differences (Sup Fig 5) and similarity among the spatial distributions of identified groups.

### Cortical layer and purified layer astrocyte RNA-Seq

#### i. Cortical layer dissection

*Aldh1L1-GFP*+ mice were transcardially perfused at P14 with ice-cold Hanks Balanced Solution (HBSS) to wash away the blood. Brains were dissected and cortical hemispheres were cut sagitally on a vibratome in ice-cold HBSS to 300 μm thick sections. Sections from 8 littermate pups were pooled for each experiment (n=3 biological replicates) and microdissected to separate upper (L2-4) and deep (L5-6) cortical layers. The L4 of the somatosensory barrel cortex, which appears as dark barrels separated by light septa under bright-field illumination, was used as an anatomical landmark for layer microdissections (Sup Fig 8). To prevent contamination with white matter astrocytes, the most superficial layers that contain pial and L1 astrocytes, and the deep subcortical white matter that contains fibrous astrocytes were discarded. For each experiment, the dissections were completed under 90 min and the tissue was kept in ice cold HBSS.

#### ii. Flow cytometry

To purify cortical layer astrocytes, tissue dissociation was performed as described previously (*17*). Briefly, cortical layer tissue were minced with a forceps and enzymatically dissociated with papain (20 U/mL) in dissociation buffer (glucose 22.5 mM, EDTA 0.5 mM, phenol red), L-cysteine (1 mM) and DNase (125 U/mL) for 80 min at 33oC. Tissue was then washed in inhibitor solution (dissociation buffer with ovomucoid (1.0 mg/mL)) and centrifuged for 5 min at 200 g. Supernatant was discarded, the tissue was resuspended in the inhibitor buffer and mechanically disrupted using a P1000 pipette. Dissociated cells were layered onto inhibitor buffer with concentrated ovomucoid (5 mg/mL) and centrifuged 5 min at 200 g. Finally, the cell pellet was resuspended in staining medium with DAPI. *Aldh1l1-GFP+* and *Aldh1l1-GFP* cells were sorted as previously described (*8*) on a BD FACS Aria II and gated on forward/side scatter, live/dead by DAPI exclusion, and GFP, using GFP and DAPI controls to set gates for each experiment (Sup Fig 8).

#### iii. RNA sequencing and analysis

Total RNA from FACS-purified cortical layer astrocytes and whole cortical layers was extracted with Trizol LS (Invitrogen) and purified using the RNeasy Kit (Qiagen). cDNA was generated from full-length RNA using the NuGEN RNA-Seq V2 kit that employs the single primer isothermal amplification method to deplete ribosomal RNA, and sheared by Covaris to yield uniform size fragments. RNASeq libraries were generated using the NuGen Ultralow kit for adapters, barcoding, and amplification and purified using the Agencourt XP magnetic beads, quality controlled with an Agilent bioanalyzer, and quantified by qPCR.

Five libraries were pooled per lane across three lanes for single end (SE50) sequencing on a HiSeq 4000. Read quality was assessed using FastQC (version 0.11.4) and 5 nucleotides at the 5’ end were trimmed. 45 nucleotide long reads were aligned to the mouse reference genome (Ensembl GRCm38) using TopHat2 (version 2.0.11) (*38*). The multiple hit parameter was (-g) was set to 1 to exclude reads with multiple genomic alignments. On average, 68 million reads were uniquely mapped to each sample (range 59-87M). Read counts per gene were calculated using SAMtoools (version 0.1.19) (*39*) and HTSeq (version 0.6.1p1) with default parameters (*40*). DESeq2 (*41*) was used to detect differentially expressed genes amongst upper and deep layer astrocytes (n=3 biological replicates) and whole cortical layers (n=2 replicates). Purified deep layer astrocytes showed low levels of contaminating oligodendrocyte marker gene expression (e.g. MBP); these genes were excluded from analysis using a mild astrocyte-specific expression threshold (astrocyte vs whole layer expression > 0.1). To identify the top differentially expressed genes between upper and deep gray matter astrocytes, an expression threshold of 5 FPKM was used with a false-discovery rate (FDR) < 0.05. The resulting list of 159 differentially expressed layer astrocyte genes is provided in Supplementary Table 9.

#### Manual immunohistochemistry

Cryosections from *Aldh1L1-GFP* mice were manually stained for GFP, astrocyte and neuron marker antibodies. Samples were subjected to heat induced antigen retrieval in 10mM sodium citrate (pH 6) buffer for 5 min at 75oC, then permeabilized and blocked in 10% goat serum in 1X PBS with 0.1% Triton X-100 (PBST) for 1 h. Primary antibodies were diluted in the blocking solution and incubated O/N at 4°C. After multiple PBST washes, samples were incubated in secondary antibodies and DAPI diluted in blocking solution for 1 h. After PBS and dH_2_O washes, samples were mounted using Fluoromount-G (SouthernBiotech). The following primary antibodies were used: chicken GFP (GFP-1020, Aves, 1:2000), mouse NEUN (MAB377, Millipore, 1:500) and rabbit GS (G2781, Sigma, 1:2000). Goat secondary antibodies conjugated to Alexa fluorophores (Molecular Probes) were used for labeling. The *Aldh1L1-GFP* samples were imaged on a Leica TCS SPE laser confocal microscope with a 40x oil immersion objective.

---

**Supplementary Figure 1:**
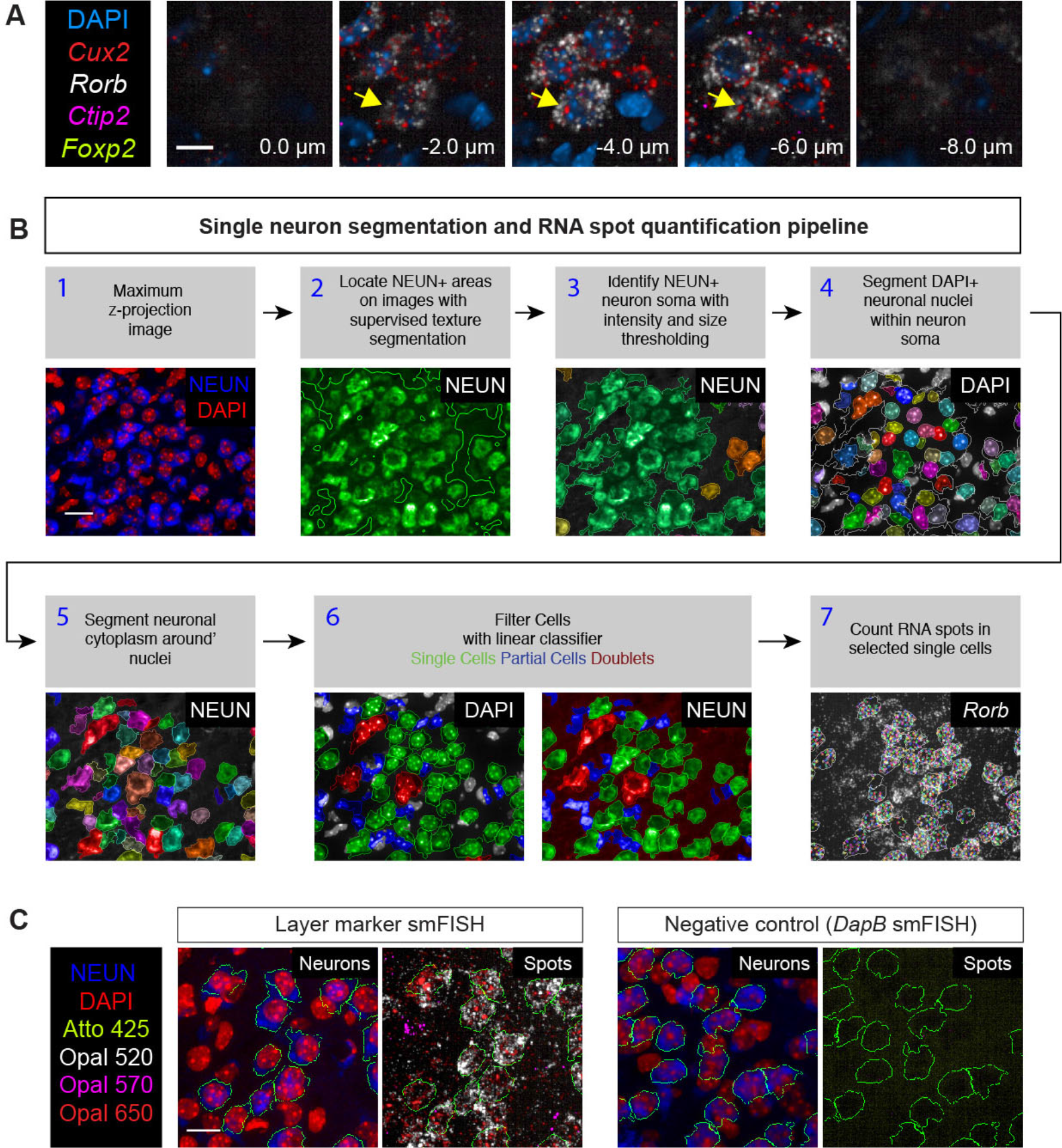
Single neuron image analysis and gene expression quantification pipeline. **A)** Individual 40X z-planes throughout *Rorb*^+^ L4 neurons. Arrow indicates a single neuron across multiple z-positions. Nuclei are marked by DAPI. For neuronal segmentation, the z-stack is collapsed into a single plane via a maximum intensity projection. **B)** Segmentation of NEUN+ neurons and quantification of gene expression in single neurons. **C)** RNAScope smFISH assay shows high signal-to-noise ratio. The background signal is assessed by comparing smFISH against layer neuron markers to bacterial *DapB* transcript negative control (targeted with four different probes in different channels). *DapB* smFISH shows little to no signal on mouse tissue sections, as expected. Quantification shown in Supplementary Figure 2. Scalebars: (A) 10 μm, (B,C) 20 μm

**Supplementary Figure 2:**
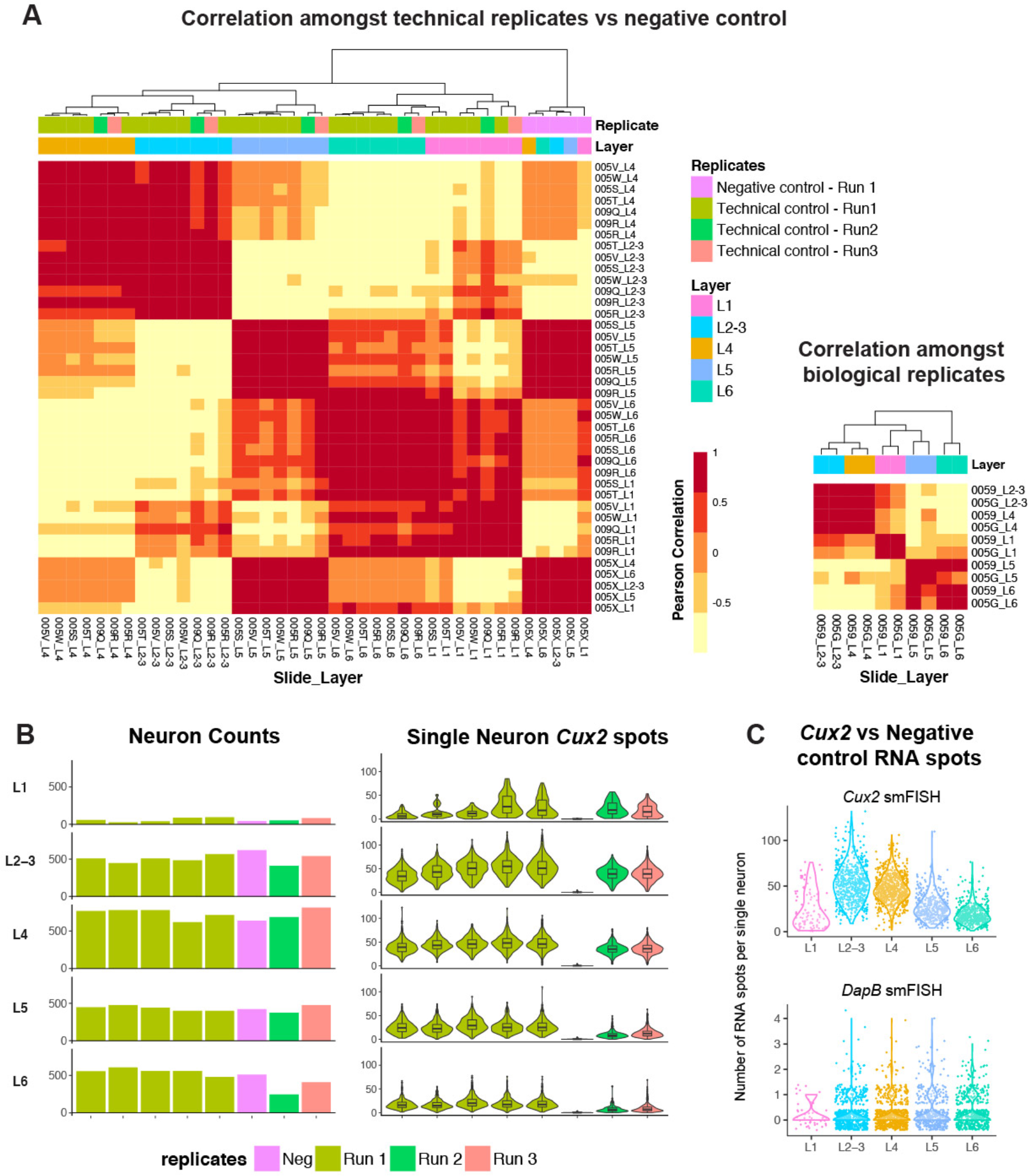
Reproducibility of single neuron gene expression measurements. **A)** Neuronal gene expression measurements are highly consistent across technical and biological replicates. (Left) Heatmap showing the pearson correlation values across expression profiles of technical replicates and the negative control. Technical replicates are consecutive P14 brain sections on different slides assayed for smFISH against 4 layer markers. Negative control was assayed for smFISH against bacterial *DapB* transcripts. To assess technical variation within a staining run, multiple replicates were assayed simultaneously on the BOND RX (Run 1). To assess batch effects, replicates were assayed on different days using different consumable reagent kits (Runs 2 and 3). To calculate the expression profiles of replicates per cortical layer, single neuron RNA spot counts for 4 layer markers (*Cux2, Rorb, Ctip2, Foxp2*) are averaged and log transformed across each layer (L1, L2-3, L4, L5, L6) in the barrel cortex. The replicate-layer expression profiles were then hierarchically clustered. As expected, cortical layers clustered across technical replicates from the same staining run as well as different batches, indicating reproducibility, while negative control layers formed a distinct cluster. (Right) Heatmap showing the pearson correlation values across expression profiles of biological replicates. Biological replicates are barrel cortex sections from two littermate P14 animals. Average layer expression profiles were calculated as described above. As expected, cortical layers from biological replicates clustered together. **B)** (Left) Quantification of neurons across cortical layers in the barrel cortex amongst replicates. Similar number of neurons are detected based on NEUN IHC and DAPI staining across technical and negative replicates. (Right) Quantification of single neuron *Cux2* expression across technical and negative replicate with violin and boxplots. Negative control shows the quantification of *DapB* expression in the same probe channel used for *Cux2* (Opal 650). The range of single cell *Cux2* expression across upper layers is highly consistent amongst technical replicates from the same staining run. Slightly lower expression is observed on batch replicate controls, yet the upper layer enrichment of Cux2 is highly similar. **C)** Quantification of single neuron Cux2 vs negative control *DapB* expression across cortical layers. The background signal of RNAScope smFISH, assessed by the numbers of *DapB* spots per cell, is 0 to 2 spots per cell.

**Supplementary Figure 3:**
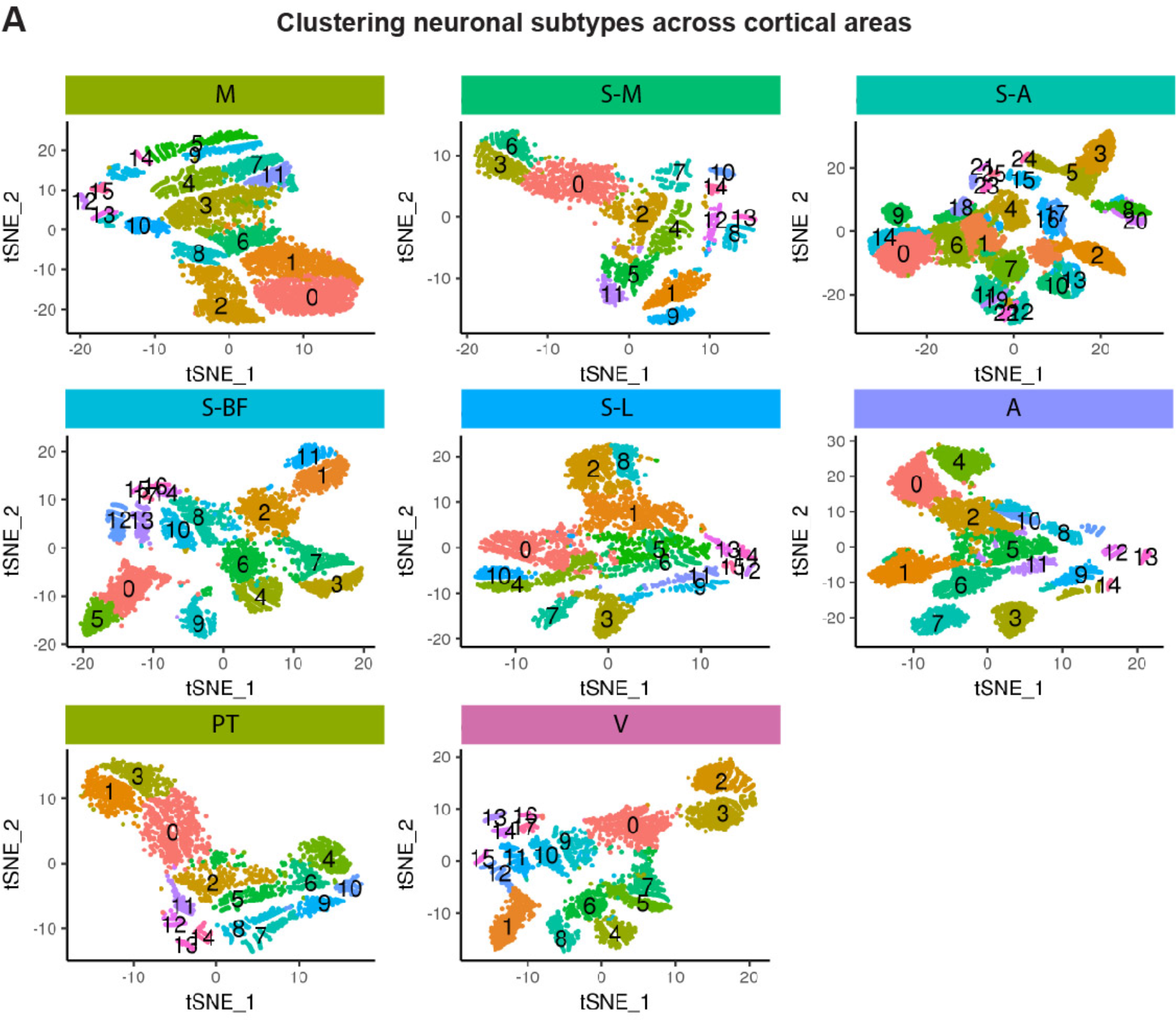
Clustering of single neurons from different cortical areas. **A)** To identify neuronal subtypes, clustering of single neuron gene expression was performed within each cortical area individually with 4 layer markers profiled. Single neuron tSNE coordinates were calculated from the expression profiles and tSNE plots were colored according to cluster assignments from above. See Supplementary Methods for details. Abbreviations: M, motor, S-A, anterior somatosensory, S-M, medial-somatosensory, S-BF, somatosensory barrel, S-L, somatosensory-lateral, PT, parietal, A, auditory, V, visual.

**Supplementary Figure 4:**
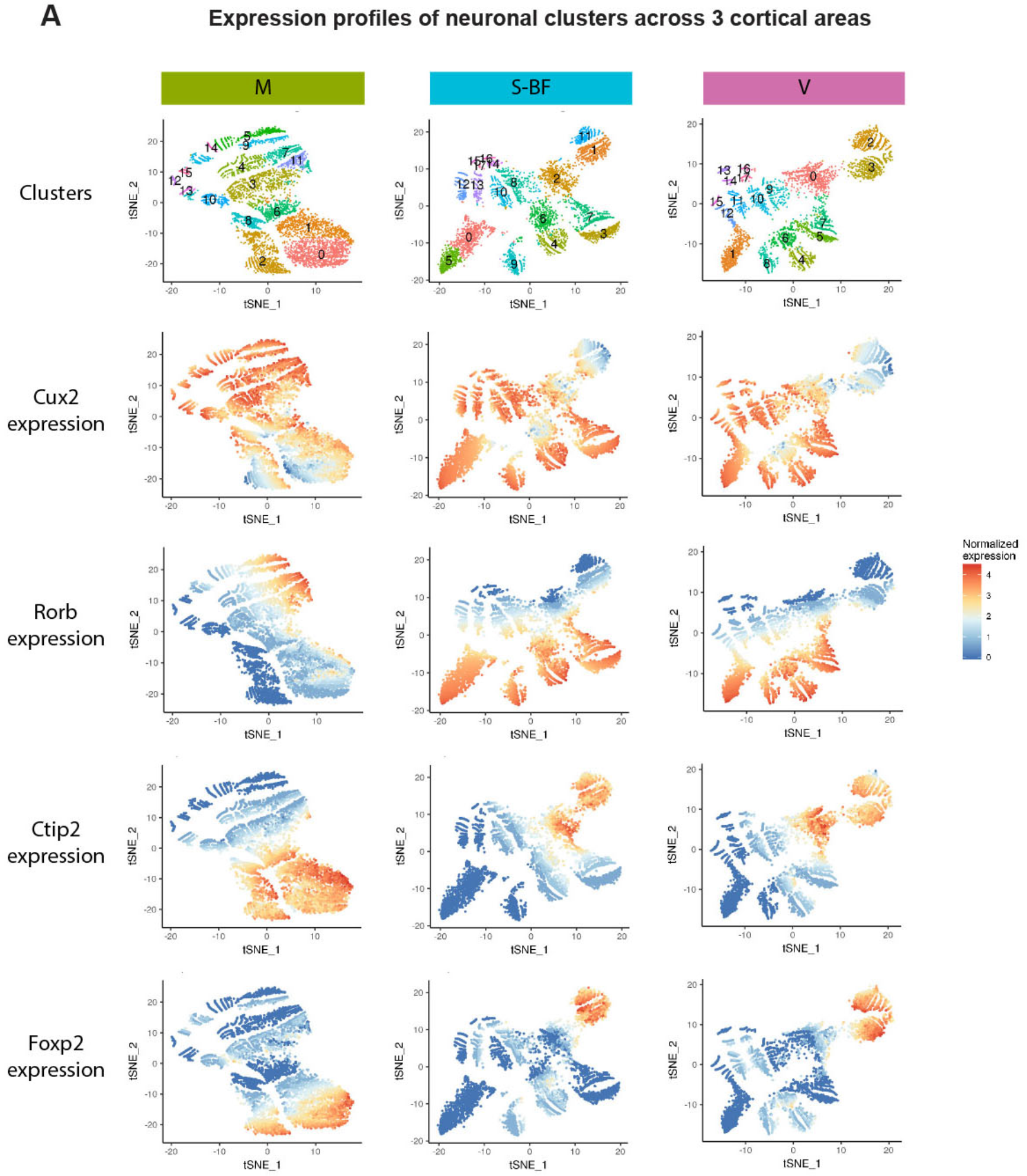
Single neuron clusters are distinguished according to layer gene expression patterns. **A)** tSNE plots across three different cortical areas (M, motor, S-BF, somatosensory barrel, V, visual) colored according to expression values.

**Supplementary Figure 5:**
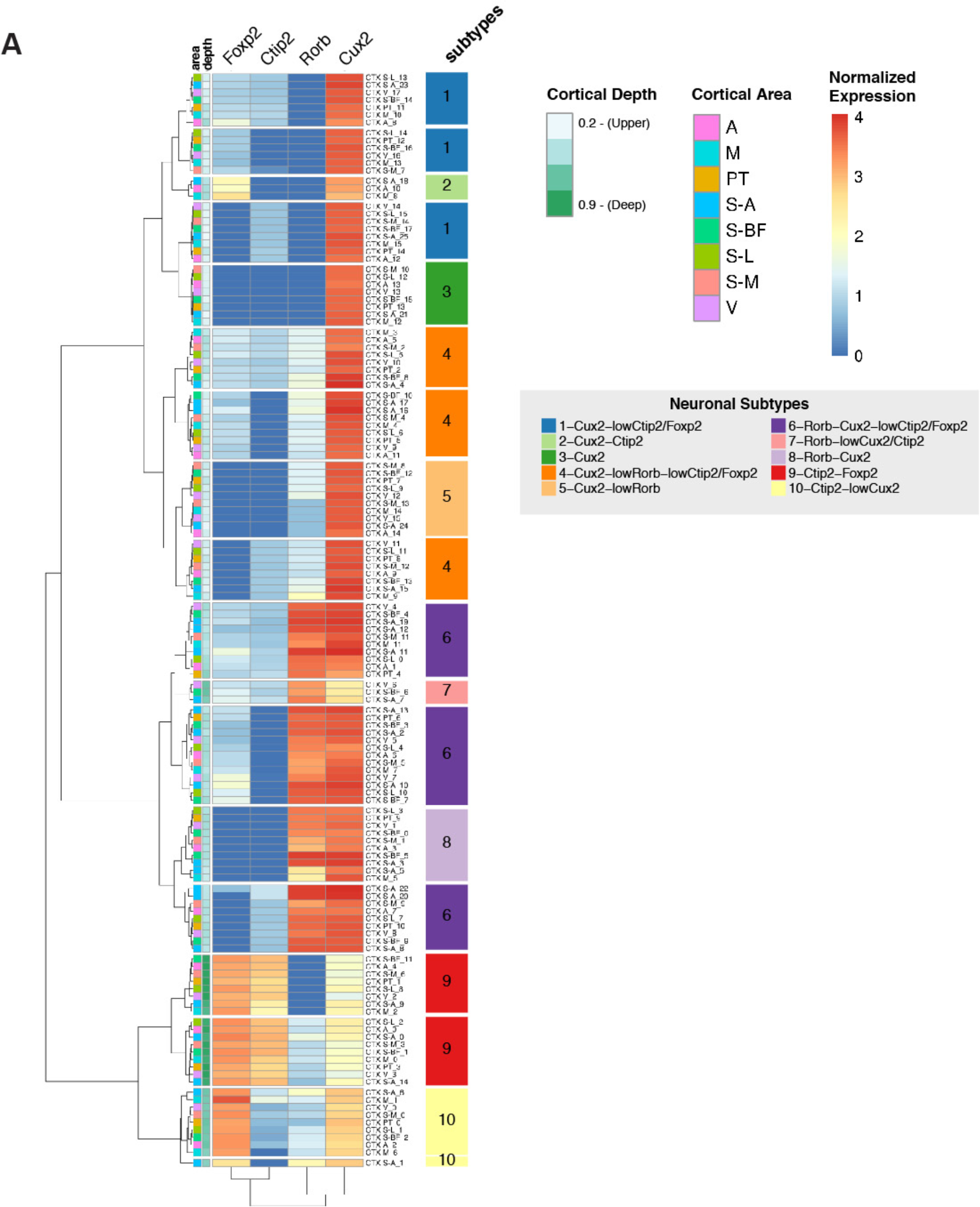
Hierarchical clustering distinguishes neuronal subtypes. **A)** Hierarchical clustering of clusters from 8 cortical areas (Supplementary Figure 3) according to mean expression profiles of each group. The clustering yielded 18 groups that were manually annotated to 10 major subtypes based on high expression differences and spatial distribution across the cortex.

**Supplementary Figure 6:**
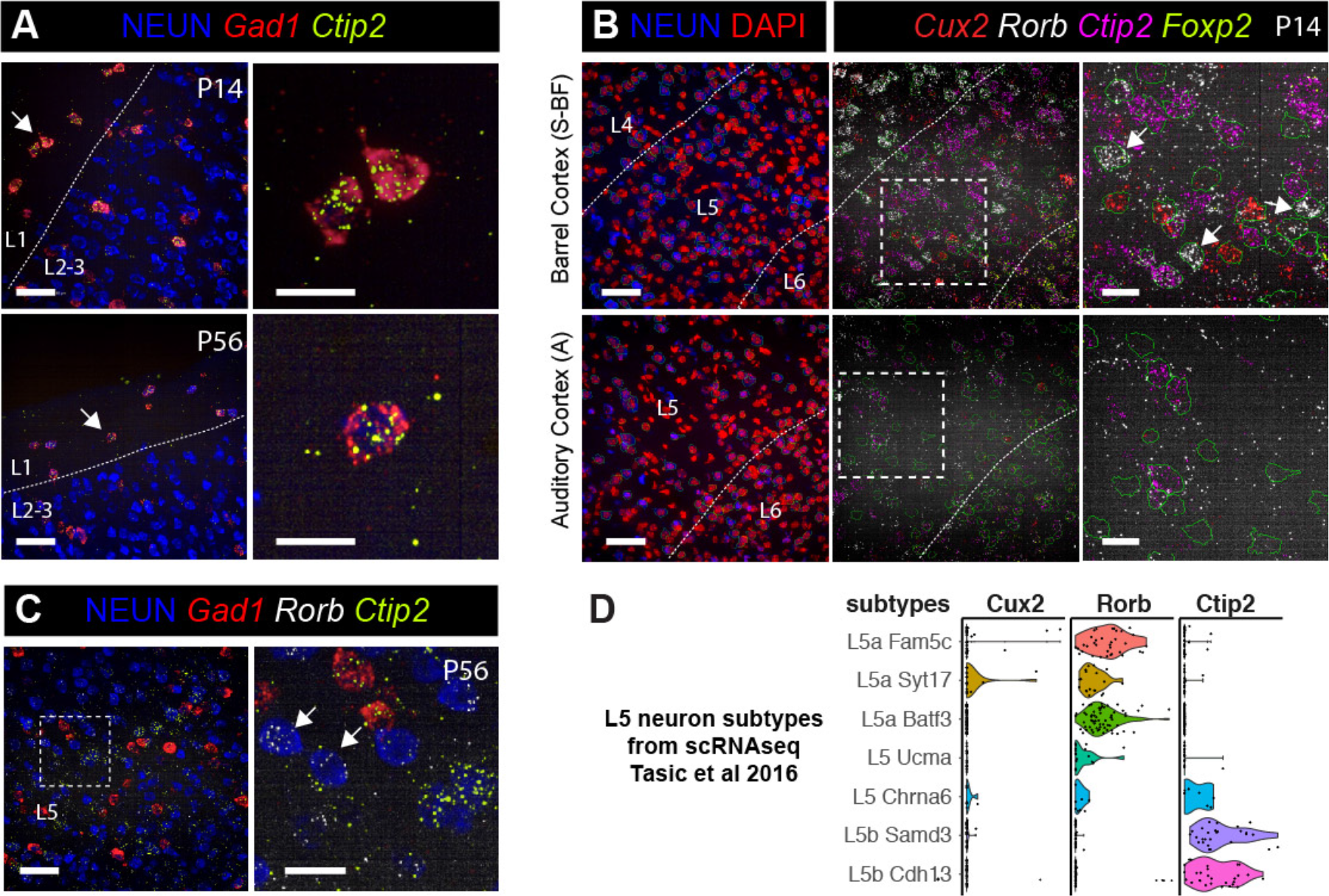
smFISH images demonstrating *Cux2*^*mid*^*Ctip2*^*mid*^-L1 and *Rorb*^*high*^*Ctip2*^*low*^-L5 neuronal populations. **A)** Neurons that co-express Cux2 and Ctip2 (cluster #2, arrows) are observed in L1. These interneurons, based on high *Gad1* expression, are present at P14 and are maintained into adulthood at P56. Right panels show higher magnification views of indicated neurons. **B)** Area enrichment of novel *Rorb*^+^ L5 subpopulations. *Rorb*^*high*^*Cux*^*mid*^*Ctip2*^*low*^ neurons (cluster #7, arrows) are observed in the L5 of the somatosensory barrel cortex, but are absent from the auditory cortex at P14. The higher magnification view of L5 areas outlined in dashed boxes shown on the right panels. **C)** *Rorb*^*high*^*Ctip2*^*low*^ neurons are maintained into adulthood at P14. **D)** Validation of *Rorb*^*high*^*Ctip2*^*low*^-L5 subtypes in a published single neuron transcriptomics datasets. Violin plots showing the segregation of *Rorb* and *Ctip2* expression amongst molecular subtypes of L5 neurons in the adult visual cortex identified by Tasic et al. Subtypes were named according to the nomenclature in the referenced study. Scalebars: (low magnification panels) 50 μm, (higher magnification panels) 20 μm

**Supplementary Figure 7:**
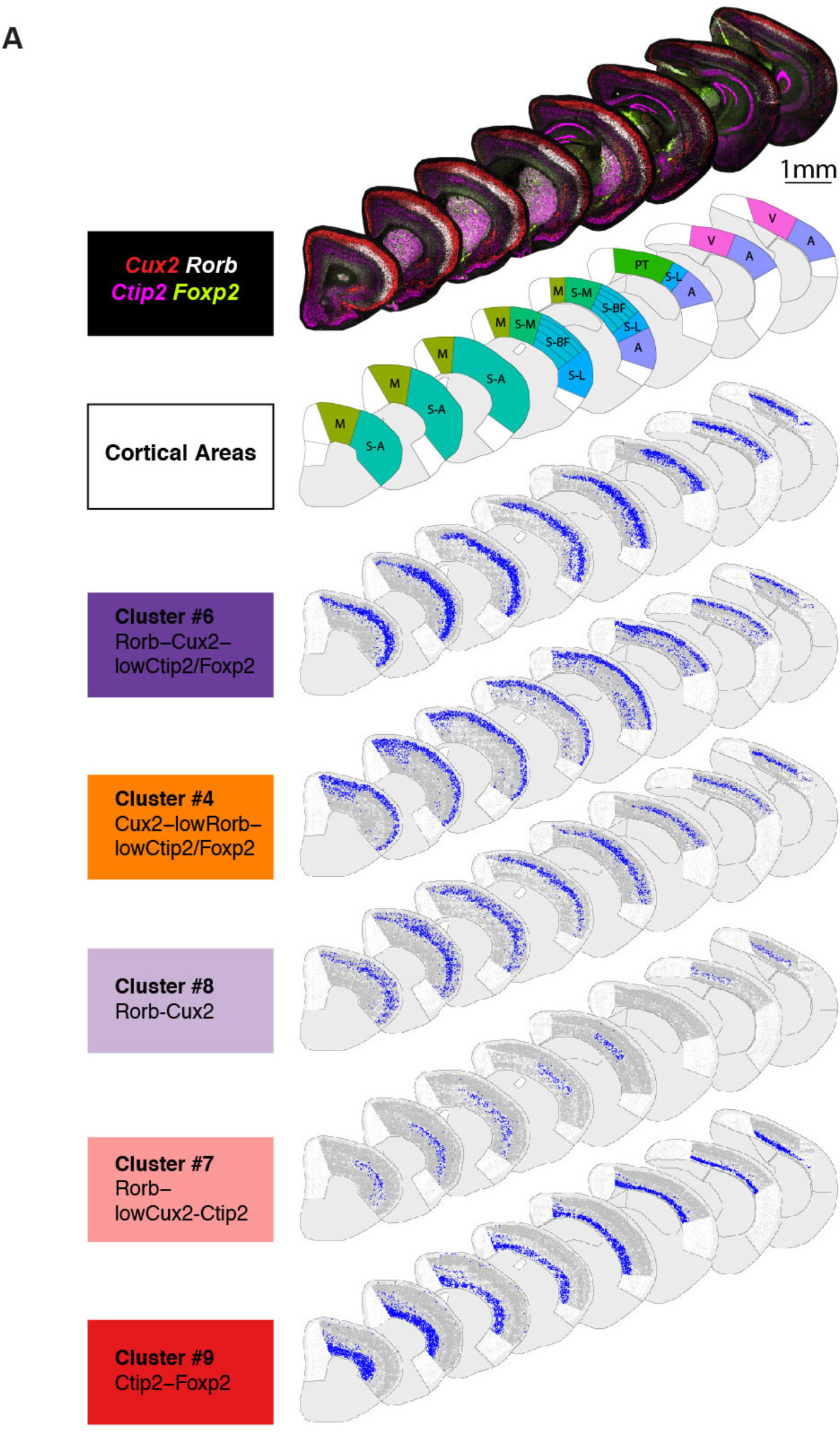
Maps showing the single cell level distribution of select neuronal subtypes. **A)** (First row) Low magnification images of P14 hemisections from eight select anatomical levels assayed for neuronal layer marker smFISH. (Second row) Maps of broad cortical areas included in neuronal subtype analysis. (Bottom rows) Maps showing the spatial distribution of individual neuronal subtype clusters. Scalebar: 1 mm

**Supplementary Figure 8:**
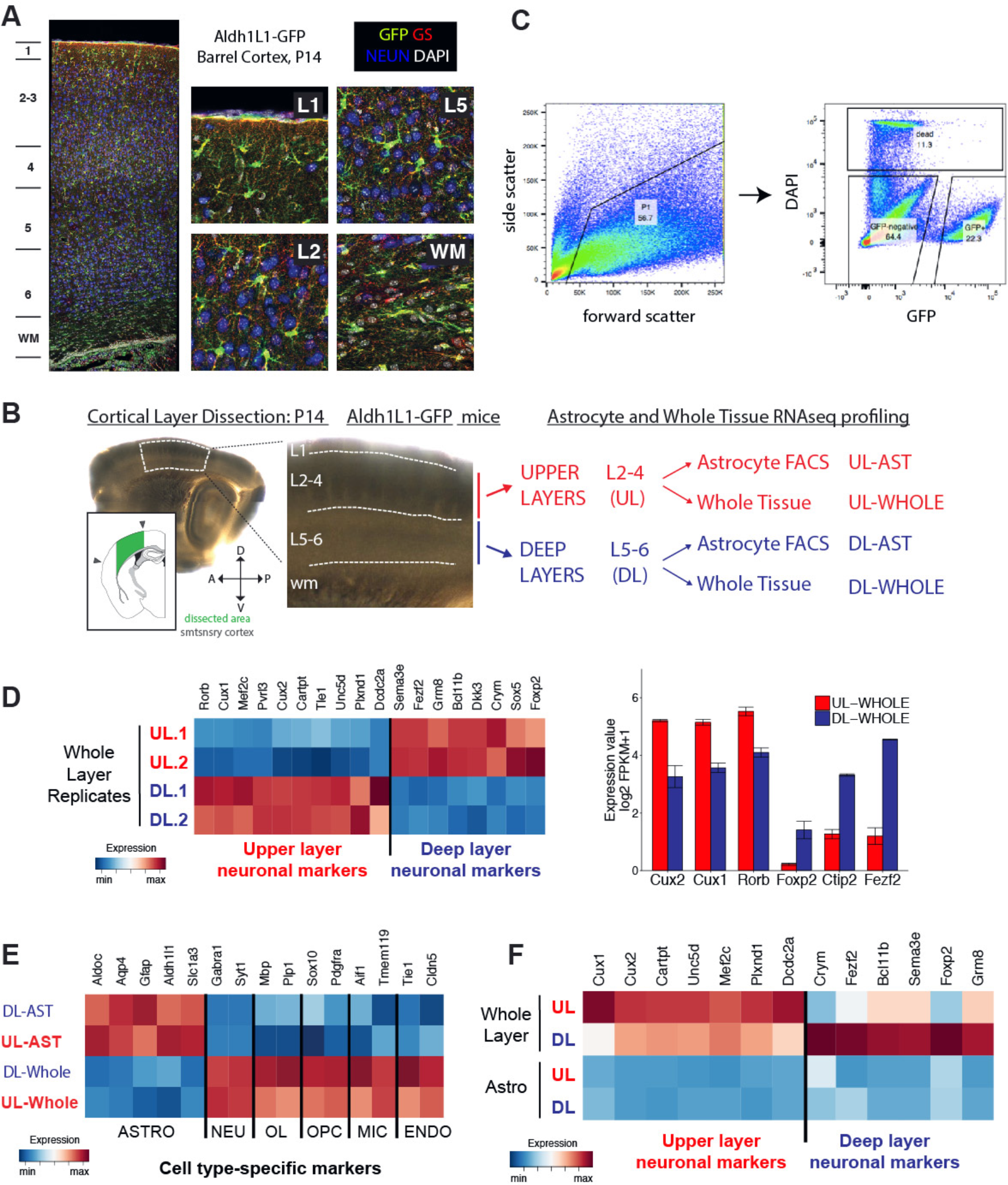
Purification and RNAseq expression profiling of upper and deep layer astrocytes. **A)** *Aldh11L1-GFP* labeling marks astrocytes across cortical layers and excludes neurons. Confocal images of antibody staining against GFP, NEUN (neuronal marker) and Glutamine Synthetase (GS, astrocyte marker) in the barrel cortex at P14. *Aldh1L1-GFP* labeling marks astrocytes throughout cortical gray matter, white matter and L1-subpia. **B)** Schematic summarizing layer astrocyte purification and gene expression profiling. (Left) Bright-field images of a sagittal P14 mouse brain slice showing the outline of the layer microdissection in the somatosensory cortex (white dashed lines & also marked green in small diagram). L4 barrels were used as an anatomical landmark. (Right) FACS-purification and RNAseq profiling strategy. **C)** *Aldh1L1-GFP*+ astrocyte isolation by FACS using scatter gates, doublet exclusion (not shown) and sorting for GFP+ cells with dead cell exclusion by DAPI staining. **D)** RNAseq expression pattern of known layer neuron markers across whole layer tissue, shown with an expression heatmap and bar-plots, validates the layer microdissection (n=3 biological replicates). **E)** Expression pattern of cell type-specific markers confirms the successful purification of astrocytes. **F)** The expression of known neuronal layer marker genes does not distinguish layer astrocytes.

**Supplementary Figure 9:**
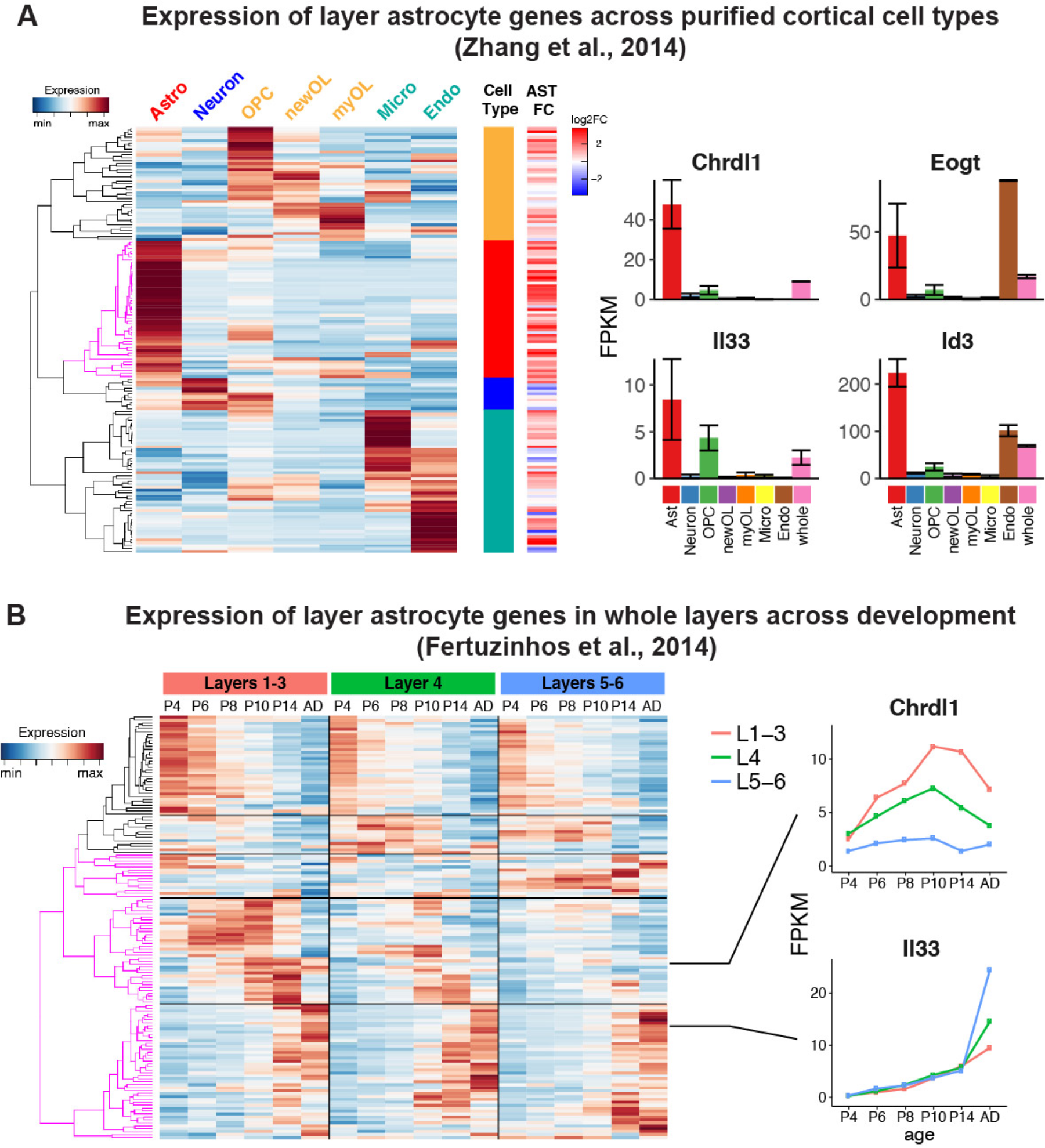
Candidate layer astrocyte genes express in laminar and astrocyte enriched manner across published cortical transcriptome dataset. **A)** The expression pattern of 163 genes differentially expressed across upper and deep layer astrocytes across purified cortical cell types. Zhang et al performed RNA-seq analysis of purified mouse cortical astrocytes, neurons, oligodendrocyte precursor cells (OPCs), newly differentiated oligodendrocytes (newOL), myelinating oligodendrocytes (myOL), microglia, and endothelial cells {Zhang:2014bt}. (Left) Heatmap shows that many candidate layer astrocyte genes show expression in astrocytes in Zhang et al’s dataset. Many genes have enriched expression in astrocytes, others also express in additional cell types. (Right) Bar plots showing expression of select candidate layer astrocyte genes across cell types. *Chrdl1* expression is highly enriched in astrocytes while *Eogt* and *Id3* also show expression on endothelial cells (validated by smFISH, data not shown). **B)** The expression of 163 candidate layer astrocyte genes across whole cortical layer tissue throughout postnatal development and adulthood. Fertuzinhos et al performed RNA-seq analysis manually dissected upper (L1-3), mid (L4) and deep (L5-6) cortical layers at different timepoints during postnatal life and adulthood {Fertuzinhos:2014fu}. (Left) Heatmap shows that many candidate layer astrocyte genes show laminar and developmentally regulated gene expression in Fertuzinhos et al’s dataset. Many genes are upregulated during early postnatal life, consistent with the commencement and progression of cortical astrogenesis after birth {Ge:2012bg} (clusters marked in magenta on the dendrogram). (Right) Most layer astrocyte candidate genes show temporally regulated expression throughput postnatal life. *Chrdl1* expression peaks during the second postnatal week yet persists into adulthood. *Il33* expression increases into adulthood.

**Supplementary Figure 10:**
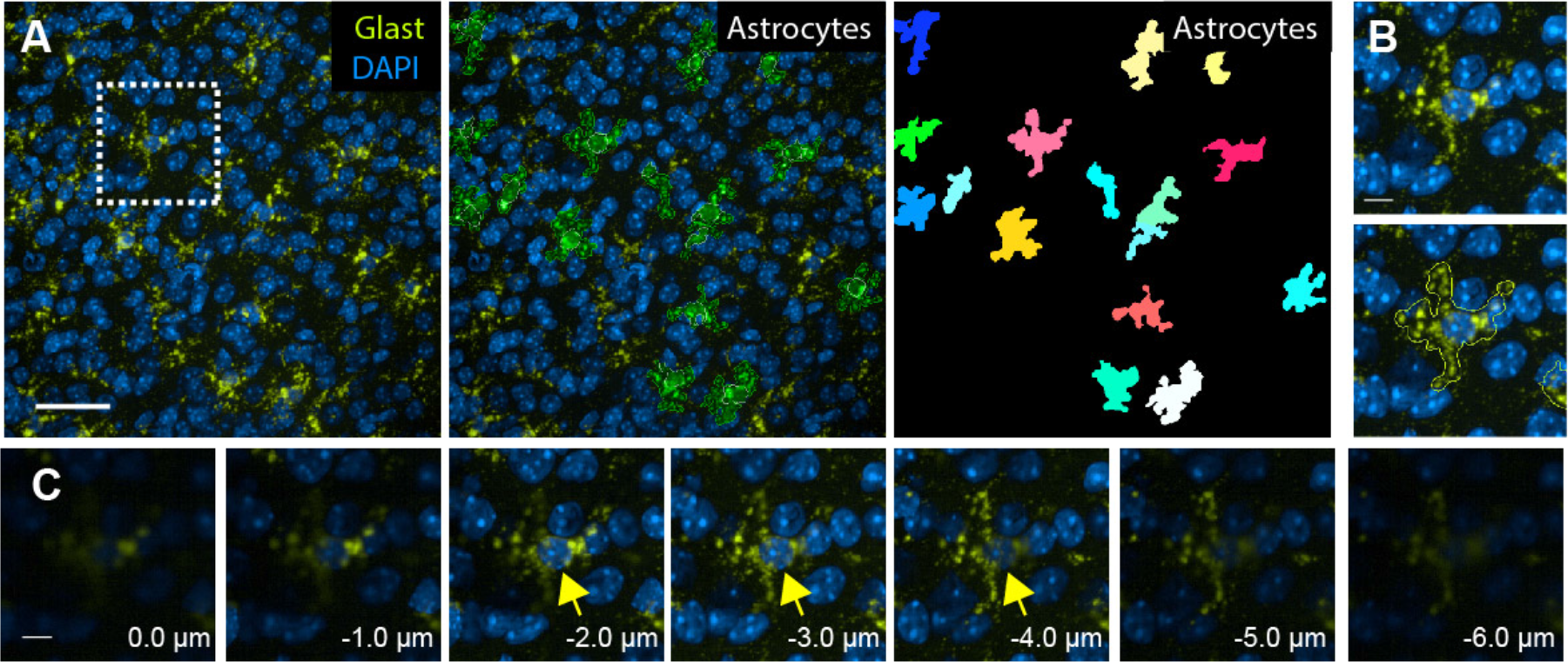
Identification of cortical astrocytes with *Glast* smFISH. **A)** (Left) Maximum z-projection image showing astrocytes in the P14 barrel cortex upper layers. (Middle) Segmentation of single astrocytes, outlined are astrocyte cell areas (green) and nuclei (white). (Right) Segmentation masks of individual astrocytes. **B)** Higher magnification image of an astrocyte indicated with dashed box in A. Bottom panel also shows the outline of the astrocyte cell area in dashed lines. **C)** Individual 40X z-planes throughout the same astrocyte. The arrow indicates the astrocyte nuclei marked by Glast and DAPI. For astrocyte segmentation, the z-stack is collapsed into a single plane via a maximum intensity projection. Scalebars: (A) 50 μm, (B,C) 10 μm

**Supplementary Figure 11:**
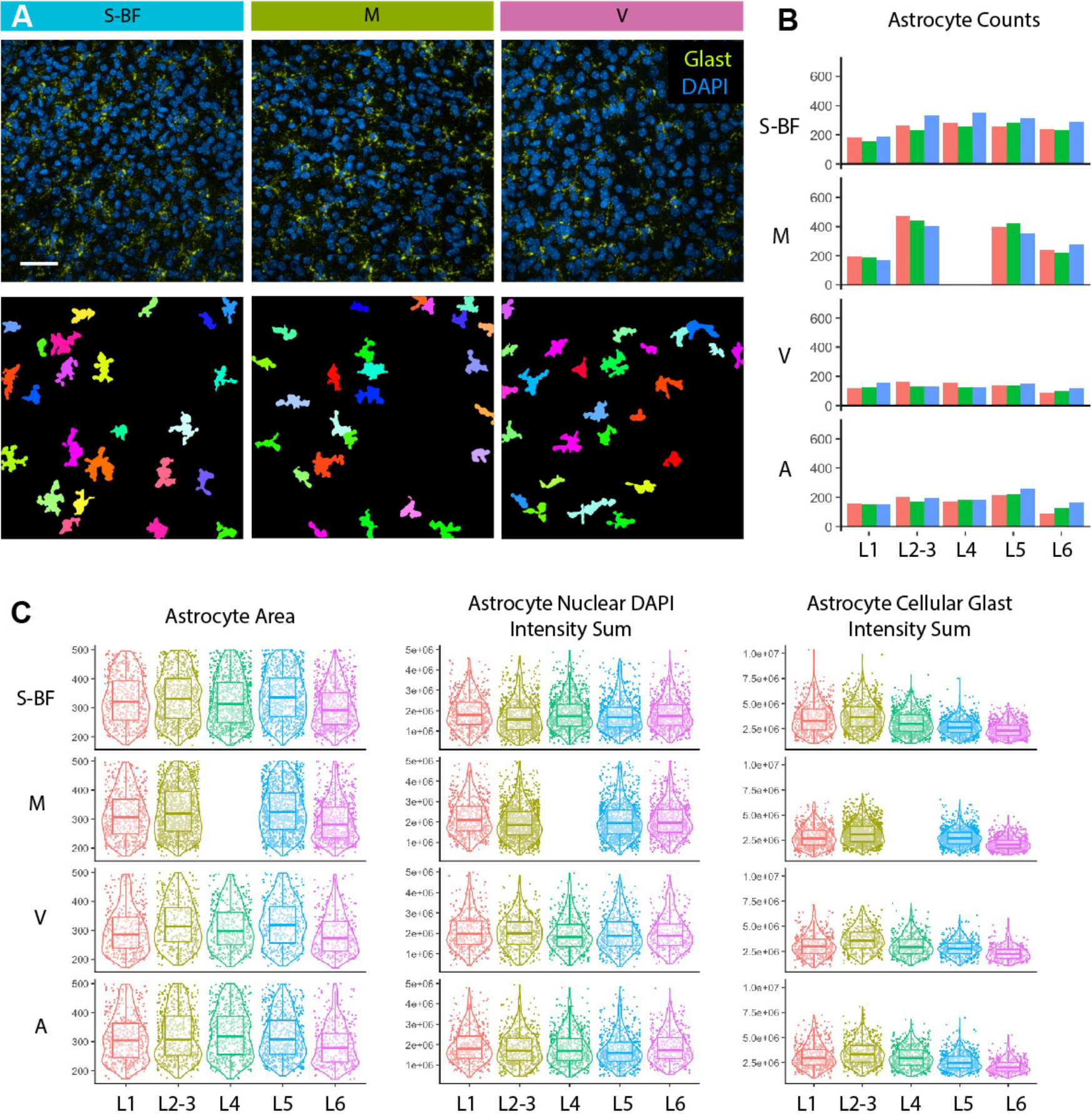
Identification of astrocytes across different cortical layers. **A)** Images (top) and segmentation masks (bottom) of astrocytes from barrel, motor and visual cortex. Midcortical layers (L4-5) are shown. **B)** Astrocyte cell counts across cortical layers and areas are consistent across three technical replicates (different colors). **C)** Astrocyte segmentation performs consistently across cortical areas. Violin, box and dot plots showing the cellular features of single astrocytes measured across four cortical areas. Deep layer astrocytes are slightly smaller and show lower expression of Glast than upper layer astrocytes. Scalebar: (A) 50 μm

**Supplementary Figure 12:**
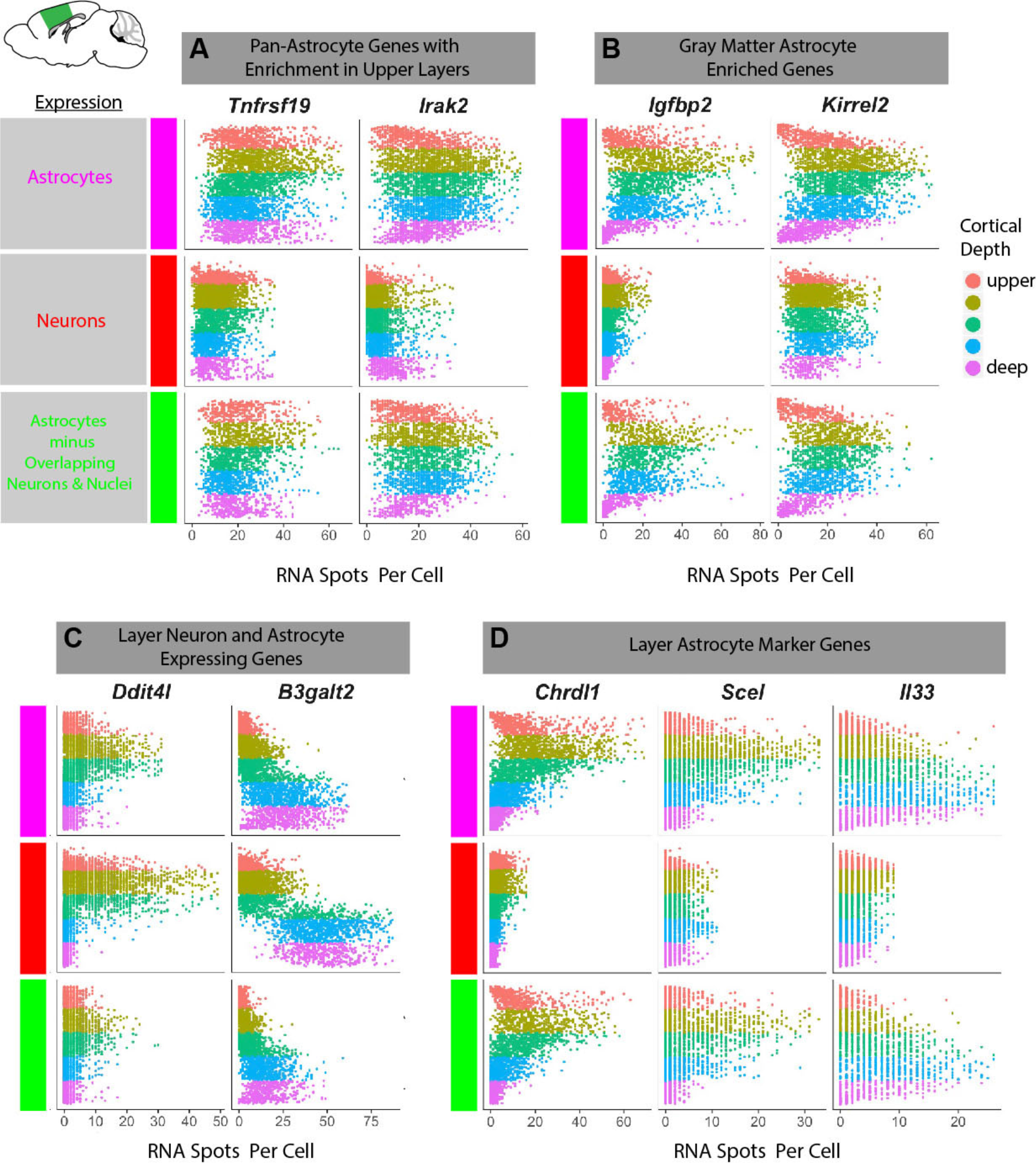
Screening and selection of top layer astrocyte markers. Quantification of single astrocyte, neuron and filtered astrocyte (i.e. removal of z-overlapping neurons and non-astrocyte nuclei) in situ expression of candidate layer astrocyte genes identifies several spatial and cell type-specific expression patterns. Screened genes show pan-astrocyte (A), gray matter astrocyte (B), astrocyte and neuron (C), and layer astrocyte enriched (D) expression patterns.

**Supplementary Figure 13:**
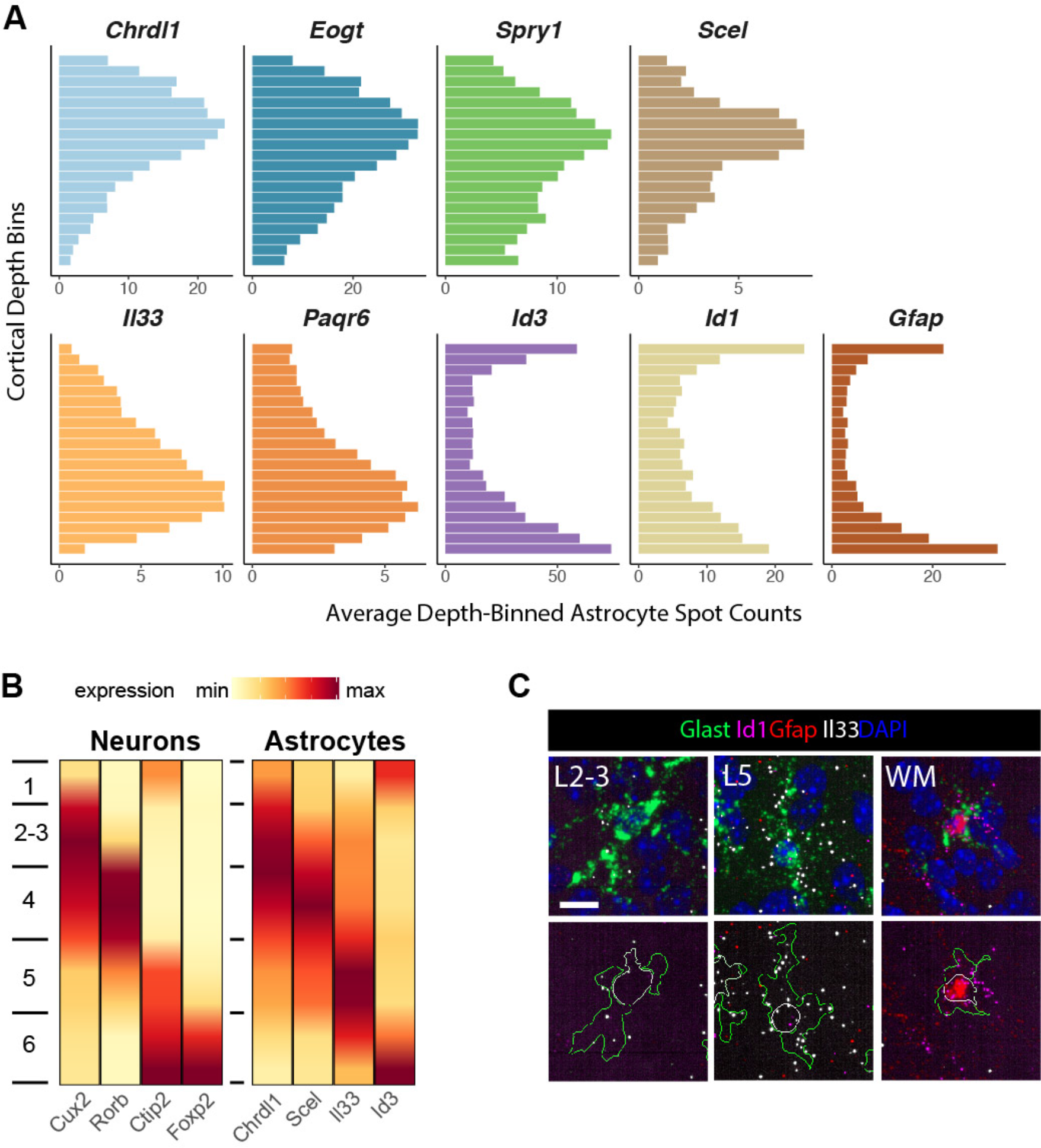
Astrocyte layer gene expression diverges from neuronal laminae. **A)** Quantification of astrocyte layer marker expression across cortical depth. Plots show the single astrocyte expression averaged across ten cortical depth bins in the P14 somatosensory cortex (n=2 pooled biological replicates). **B)** Interpolated tile expression plots comparing neuron vs astrocyte layer marker expression across cortical depth in the P14 barrel cortex (n=3 pooled tissue sections from one biological replicate). Astrocyte layer expression domains diverge from sharply refined neuronal laminae. **C)** *Il33* expression is enriched in L5 astrocytes but absent from white matter astrocytes at P14. Scalebar: (C) 10 μm

**Supplementary Figure 14:**
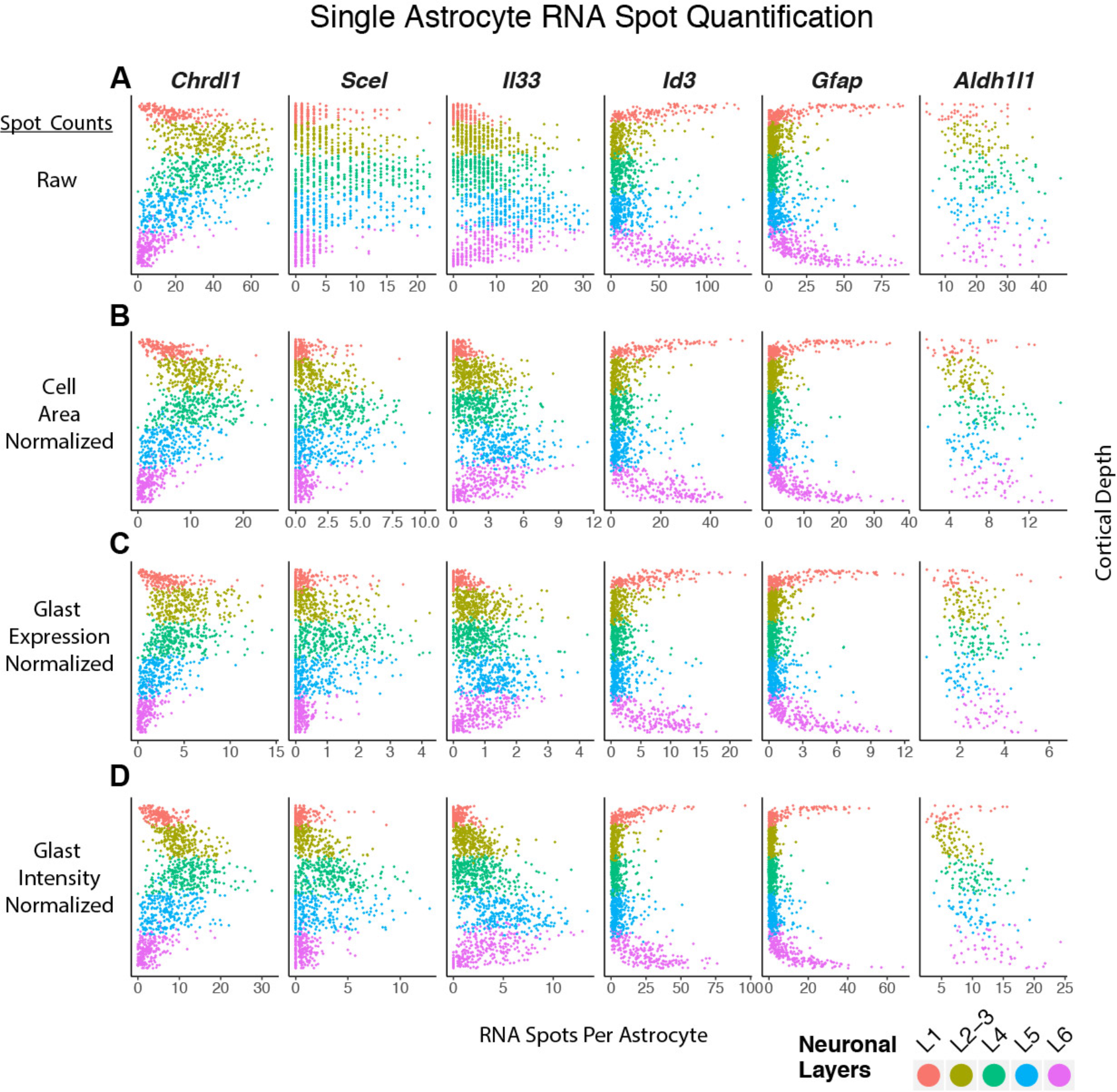
The observed astrocyte layer gene expression patterns are not artifacts of cell size or *Glast* expression level differences. Quantification of single astrocyte expression across cortical depth in the P14 barrel cortex (n=3 pooled tissue sections from one biological replicate). The expression of identified layer astrocyte markers, the white matter astrocyte marker *Gfap* and the pan-astrocyte marker *Aldh1l1* are plotted as single cell RNA spot counts that are (A) raw, or normalized to (B) astrocyte area, (C) single cell *Glast* spot counts, and (D) single cell *Glast* signal intensity.

**Supplementary Figure 15:**
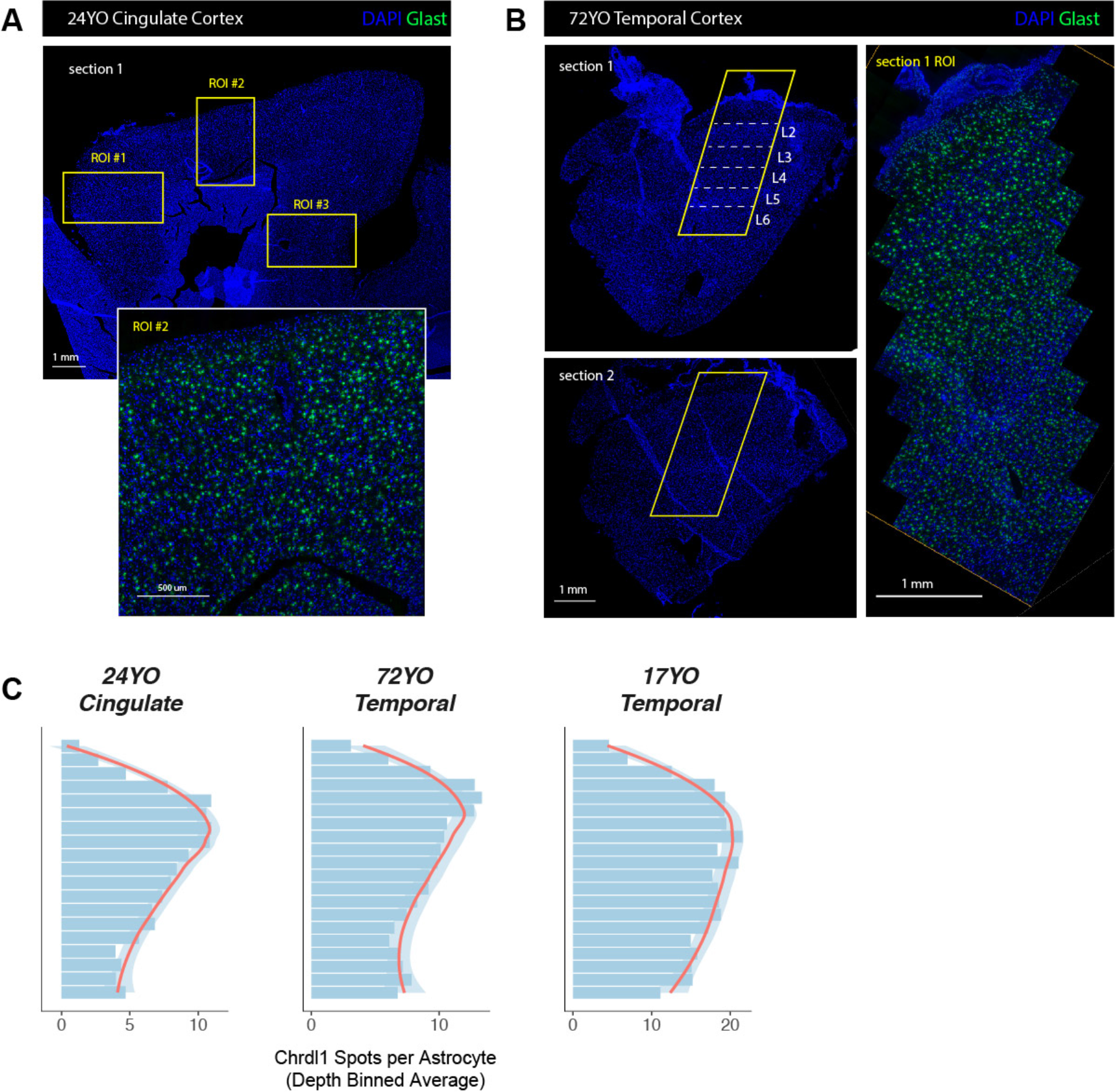
*Chrdl1* expression is enriched in upper layer astrocytes in the adult human cortex. **A)** Astrocytes in the 24 year old cingulate cortex. Low magnification images show DAPI staining of a section through the cingulate cortex (top) and Glast smFISH of the ROI #2 (bottom). Boxed regions of interest were imaged at 40X to quantify layer astrocyte expression of *Chrdl1*. **B)** Astrocytes in the 72 year old temporal cortex. Low magnification images show DAPI staining of two sections through the temporal cortex (left) and Glast smFISH of the ROI on the first section (right). Boxed regions of interest were imaged at 40X to quantify layer astrocyte expression of *Chrdl1*. **C)** Quantification of depth binned average single astrocyte expression of Chrdl1 across the 24YO cingulate, 72YO temporal and 17YO temporal cortex samples. Scalebars: (DAPI) 1 mm, (DAPI-*Glast*) 0.5 mm

**Supplementary Figure 16:**
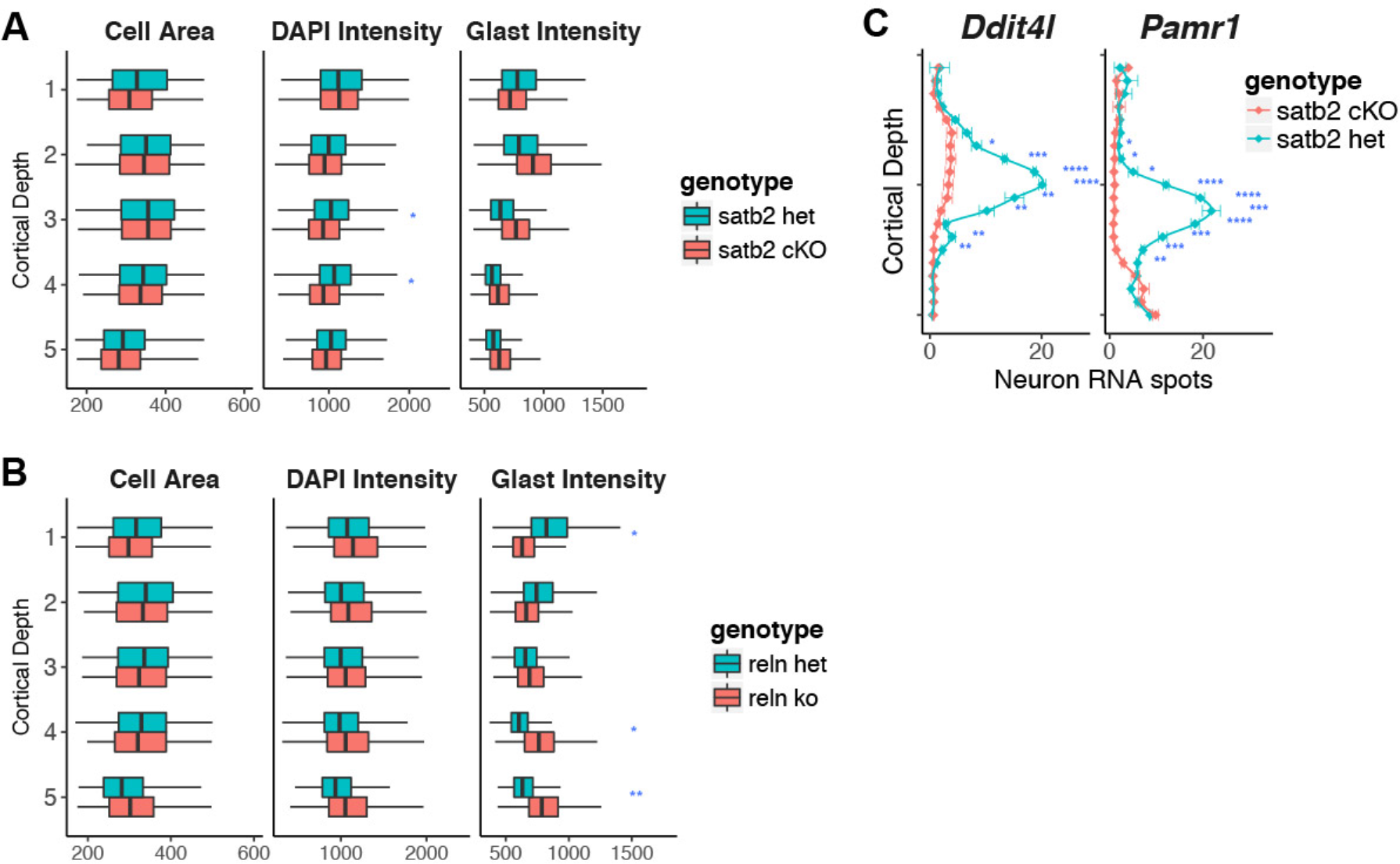
Astrocyte cellular phenotypes in neuronal layer switch experiments and loss of L4 neuron gene expression in *Satb2* cKO. **A)** Boxplots comparing astrocyte area, DAPI and Glast intensity between *Satb2* cKO and littermate controls in the P14 barrel cortex. No significant change is observed in these astrocyte features upon *Satb2* cKO (n=3 pooled biological replicates per genotype). **B)** Boxplots comparing astrocyte area, DAPI and Glast intensity between *Reeler* and littermate controls in the P14 barrel cortex. In *Reeler*, the difference in *Glast* expression between superficial and deep astrocytes is inversed, consistent with the inversion of astrocyte layers based on marker gene expression (n=3 pooled biological replicates per genotype). **C)** Quantification of cortical depth binned neuronal layer marker expression in cKO vs control (n=3 pooled biological replicates per genotype). *Satb2* cKO shows loss of L4 neuron gene expression based on additional L4 markers, *Ddit4l* and *Pamr1*. Data represent mean ± s.d. *P < 0.05,**P < 0.01, ***P < 0.001.

**Supplementary Figure 17:**
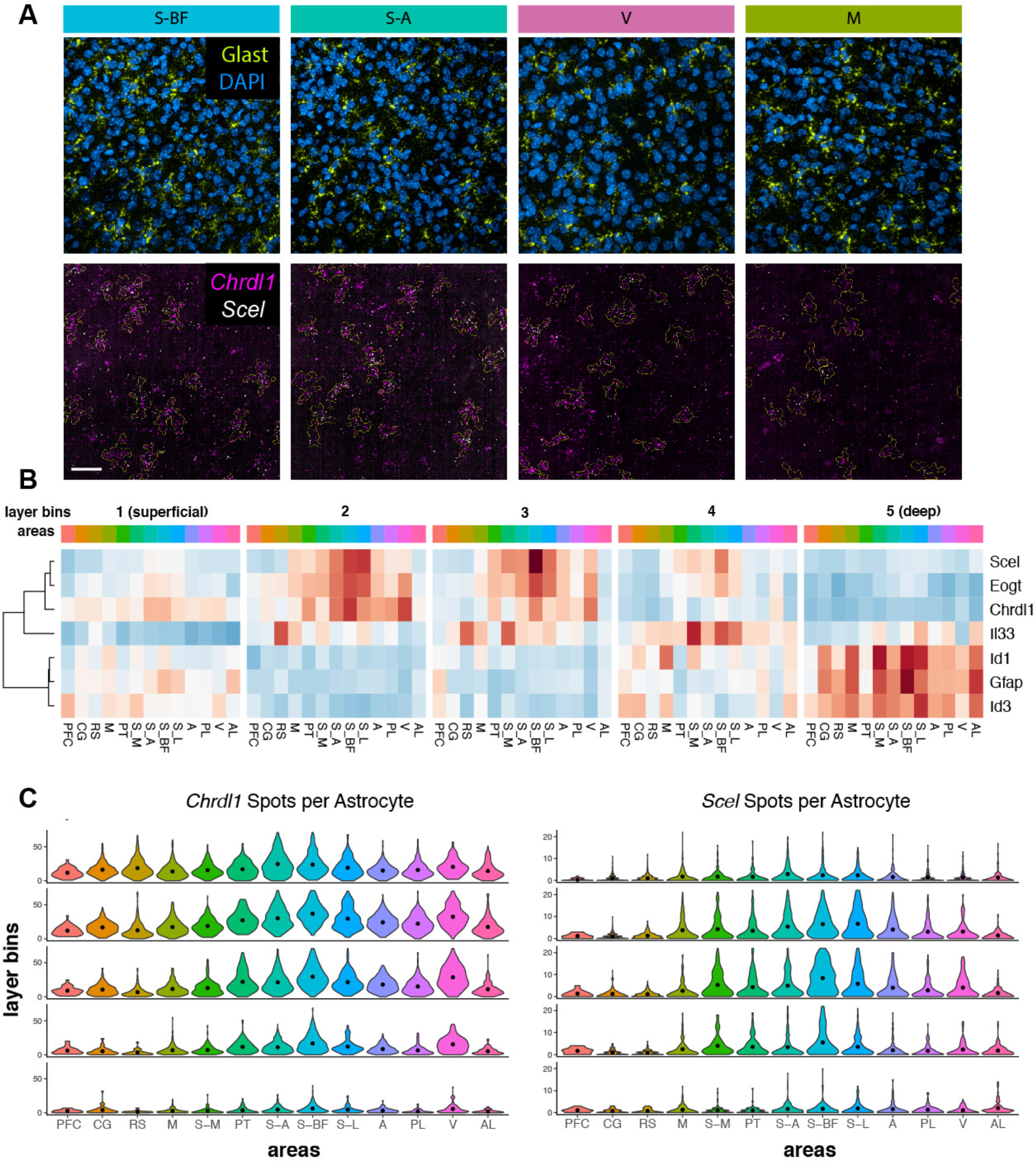
Astrocyte layer gene expression varies across cortical areas. **A)** Images showing the enrichment of *Scel* and *Chrdl1* expression in somatosensory areas over motor and visual cortex. In the bottom panels, the single astrocyte segmentations are shown in green. **B)** Expression heatmap showing the expression of layer astrocyte markers across cortical depth and areas (assayed over n=10 tissue sections from one biological replicate). **C)** Violin plots showing the quantification of single astrocyte expression of *Chrdl1* and *Scel* across cortical layers and areas.

**Supplementary Figure 18:**
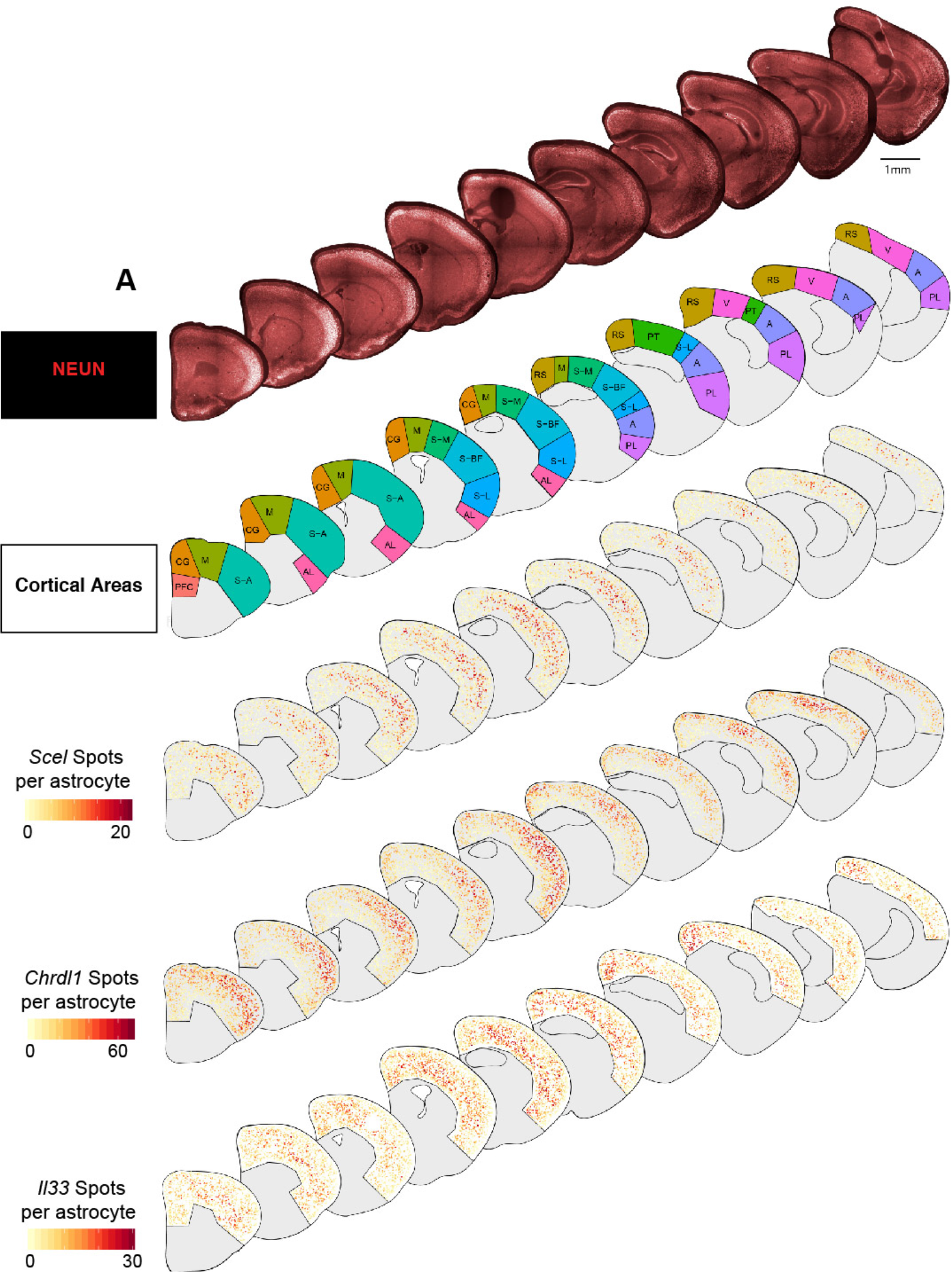
Maps showing the single cell level distribution of select neuronal subtypes. **A)** (First row) Low magnification images of P14 hemisections from ten anatomical levels assayed for NEUN IHC. (Second row) Maps of broad cortical areas included in analysis of regional astrocyte gene expression. (Bottom rows) Maps showing single astrocyte expression of *Scel*, *Chrdl1* and *Il33* across the cortex. Scalebar: 1 mm

## Supplementary Table Legends

**Supplementary Table 1: Single cortical neuron smFISH dataset.** Table listing cellular, anatomical and gene expression measurements of 69,318 single neurons identified across the P14 cortex. The cluster assignments of 46,887 single neurons used for subtype identification across 8 broad cortical areas are also listed. Every row is a single neuron and the table columns are described in the “column_metadata” sheet.

**Supplementary Table 2: Single cortical astrocyte smFISH screen dataset.** Table listing cellular, anatomical and gene expression measurements of 41,187 single astrocytes screened in the somatosensory cortex across two biological replicates. 46 candidate layer astrocyte markers as well as the pan-astrocyte marker *Aldh1l1* and the white matter astrocyte marker *Gfap* were multiplexed with the astrocyte marker *Glast* across multiple slides, these are listed under “slide_metadata”. Every row is a single astrocyte and the table columns are described in the “column_metadata” sheet.

**Supplementary Table 3: RNAscope probes used in this study.** Table listing all of the RNAScope probes, their mRNA target regions and ACD catalog numbers.

**Supplementary Table 4: Automated histology protocols and reagents.** Tables listing the automated 4-plex RNAScope smFISH and IHC protocol used on the Leica BOND RX and the consumable reagents.

**Supplementary Table 5: Imaging settings. Tables listing t**he fluorophores, light sources, exposure times and emission filters used for mouse and human tissue imaging.

**Supplementary Table 6: The single neuron segmentation and gene expression quantification pipeline.** Table listing all of the steps and settings used in the Harmony software.

**Supplementary Table 7: The single astrocyte segmentation and gene expression quantification pipeline.** Table listing all of the steps and settings used in the Harmony software.

**Supplementary Table 8: List of abbreviations for cortical areas.** Table listing all the broad cortical areas examined in this study.

**Supplementary Table 9: Cortical layer astrocyte RNAseq data.** The RNAseq expression pattern and differential gene expression statistics of 159 candidate layer astrocyte markers. The list of 46 top genes screened with smFISH is also provided. The table columns are described in the “column_metadata” sheet.

